# MyD88-Syk axis is a critical determinant for inflammatory response in macrophages

**DOI:** 10.1101/844076

**Authors:** Young-Su Yi, Han Gyung Kim, Ji Hye Kim, Woo Seok Yang, Eunji Kim, Deok Jeong, Jae Gwang Park, Nur Aziz, Suk Kim, Narayana Parameswaran, Jae Youl Cho

## Abstract

Inflammation, a vital immune response to infection and injury, is mediated by macrophage activation. Spleen tyrosine kinase (Syk) and myeloid differentiation primary response 88 (MyD88) regulate the inflammatory response in macrophages, however, their roles and the underlying mechanisms are still largely unknown. In this study, the role of the MyD88-Syk axis and the mechanism by which Syk and MyD88 cooperate during macrophage-mediated inflammatory responses were investigated. Inhibition of Syk or MyD88 decreased both generation of reactive oxygen species (ROS)/reactive nitrogen species (RNS) and consequent ROS/RNS-induced phagocytic activity in lipopolysaccharide (LPS)-stimulated RAW264.7 cells. Syk inhibition downregulated expression of ROS/RNS-generating enzymes by inhibiting the nuclear factor-kappa B (NF-κB) signaling pathway and phagocytic activity by suppressing suppressor of cytokine signaling 1 (SOCS1) via its nitration in the LPS-stimulated RAW264.7 cells. Inhibition of ROS/RNS generation suppressed SOCS1 nitration, leading to a decrease in the phagocytic activity of macrophages. Syk was activated by the interaction with MyD88, and the tyrosine 58 residue (Y58) in the hemi-immunoreceptor tyrosine-based activation motif (ITAM) of MyD88 was critical for their interaction and the consequent activation of Syk in macrophages. Src activated MyD88 by phosphorylation at Y58 via the Src kinase domain. Moreover, LPS-induced formation of filamentous actin (F-actin) and Ras-related C3 botulinum toxin substrate 1 (Rac1) activation induced Src activation. Conclusively, these results suggest that the MyD88-Syk axis plays a pivotal role in macrophage-mediated inflammatory responses by inducing ROS/RNS generation and phagocytic activity via activation of Src and its upstream stimulators, F-actin and Rac1.

## Introduction

Inflammation is an innate immune response mediated mainly by myeloid immune cells, such as macrophages, to protect the body from infection by various pathogens and cellular danger signals (Janeway & Medzhitov, 2002; Takeuchi & Akira, 2010). The inflammatory response occurs from the interaction of pattern recognition receptors (PRRs), such as Toll-like receptors (TLRs), RIG-I-like receptors (RLRs), NOD-like receptors (NLRs), and C-type lectin receptors (CLRs), with the pathogen-associated molecular patterns (PAMPs) and the danger-associated molecular patterns (DAMPs) (Janeway & Medzhitov, 2002; Takeuchi & Akira, 2010). This in turn activates the inflammatory signaling pathways, such as nuclear factor-kappa B (NF-κB), activator protein-1 (AP-1), and interferon regulatory factors (IRFs) (Byeon et al, 2012; Yang et al, 2014; Yi et al, 2014; Yu et al, 2012). The activation of these inflammatory signaling pathways is mediated by the initiation of rapid signal transduction cascades stimulating receptor-associated intracellular adaptors, such as MyD88 and TRIF, and a variety of intracellular signaling molecules, such as Src, spleen tyrosine kinase (Syk), phosphoinositide 3-kinase (PI3K), Akt, phosphoinositide-dependent kinase 1 (PDK1), inhibitor of κB (IκB) kinase α/β (IKKα/β), and IκB in the NF-κB signaling pathway; c-Jun N-terminal kinase (JNK), p38, and extracellular signal-regulated kinase (ERK) in the AP-1 signaling pathways; and TANK, TANK-binding kinase 1 (TBK1), and IKKε in the IRF signaling pathway (Byeon et al, 2012; Yang et al, 2014; Yi et al, 2014; Yu et al, 2012). As a result of the activation of these inflammatory signaling pathways, macrophages induce inflammatory responses in various ways, such as facilitating the generation of inflammatory mediators, including nitric oxide (NO) and prostaglandin E_2_ (PGE_2_); upregulating the expression of inflammatory enzymes, including inducible NO synthase (iNOS), cyclooxygenase-2 (COX-2), and matrix metalloproteinases (MMPs); and upregulating the expression of pro-inflammatory cytokines, including interleukin (IL)-1β, IL-6, tumor-necrosis factor (TNF)-α, interferon-gamma inducible protein 10 (IP-10), and interferon (IFN)-β (Byeon et al, 2012; Yang et al, 2014; Yi et al, 2014; Yu et al, 2012). Although inflammation is a protective immune response, chronic inflammation rearranges tissue architecture and consequently destroys tissues, leading to the pathogenesis of numerous inflammatory/autoimmune diseases such as rheumatoid arthritis, type I diabetes mellitus, atherosclerosis, Crohn’s diseases, systemic lupus erythematosus, Sjögren syndrome, inflammatory bowel diseases, multiple sclerosis, amyotrophic lateral sclerosis, asthma, chronic obstructive pulmonary disease, sepsis, and even cancers (Crusz & Balkwill, 2015; Todoric et al, 2016; Yi, 2016a; Yi, 2016b; Yi, 2017; Yi, 2018a; Yi, 2018b). Therefore, many efforts have been made to discover the molecular mechanisms that induce inflammatory responses, identify and validate molecular targets to suppress inflammatory responses, and develop therapeutic agents to effectively ameliorate the symptoms of inflammatory/autoimmune diseases and cancers.

Syk, a non-receptor tyrosine kinase with a molecular weight of around 72 kDa, belongs to a tyrosine kinase family considered to play pivotal roles in various biological functions. Syk has three main domains: an N-terminal Src Homology 2 (SH2-N) domain, a C-terminal Src Homology 2 (SH2-C) domain, and a kinase domain (Yi et al, 2014). An alternatively spliced form of Syk, Syk-B, has also been identified in some cells (Rowley et al, 1995), but the exact function of Syk-B and alternative splicing mechanisms are still unknown. Syk’s structure is highly conserved among species. Human and mouse Syk share 92% amino acids, and Syk-like molecules are found in other species, including rat, pig, fruit fly, and hydra (Rowley et al, 1995; Steele et al, 1999; Taniguchi et al, 1991; Ziegenfuss et al, 2008). Syk catalyzes the phosphorylation of a number of proteins at their tyrosine residues (Xue et al, 2013), and these phosphorylated proteins serve as the scaffolds to downstream effector proteins for their subsequent activation, including the signaling molecules activated in the NF-κB signaling pathway (Lee et al, 2009; Mocsai et al, 2010; Yi et al, 2014). Syk is widely expressed in hematopoietic and non-hematopoietic cells, such as immune cells, neuronal cells, epithelial cells, vascular endothelial cells, and fibroblasts (Yi et al, 2014). Syk is a critical player in diverse biological functions, including cellular adhesion, innate pathogen recognition, tissue damage recognition, antibody-dependent cellular cytotoxicity, bone metabolism, platelet functions, and vascular development (Mocsai et al, 2010; Yi et al, 2014).

Myeloid differentiation primary response 88 (MyD88), an intracellular universal adaptor of TLRs and IL-1 receptors (IL-1Rs), stimulates the transduction cascades of inflammatory signaling pathways. MyD88 is ubiquitously expressed in many tissues and consists of three main domains: an N-terminal death domain (DD), an intermediate domain (ID), and a C-terminal Toll-interleukin-1 receptor (TIR) domain (Deguine & Barton, 2014). The TIR domain interacts with other TIR domain-containing proteins, while the DD interacts with IL-1R-associated kinase 1 and 4 (IRAK1 and IRAK4) through homotypic DD interaction. MyD88s, a splicing variant of MyD88 which lacks the ID, is also found (Janssens et al, 2002), but unlike MyD88, MyD88s is not able to activate the inflammatory NF-κB signaling pathway (Janssens et al, 2003). The interaction between MyD88 and IRAK1/4 induces IRAK4-mediated IRAK1 phosphorylation which in turn triggers the activation of other signal transduction cascades of inflammatory signaling pathways such as NF-κB and AP-1 (Cao et al, 1996). A large number of studies have demonstrated that MyD88 plays a central role in the innate immune response (Oda & Kitano, 2006). MyD88 is also involved in other biological functions, including carcinogenesis, autophagy, immune activation, learning activity, anxiety, and motor functions (Cuda et al, 2014; Holtorf et al, 2018; Lim et al, 2012; Mohs et al, 2019; Siracusano et al, 2016; van der Vaart et al, 2014).

Despite a number of previous studies of Syk and MyD88, their roles in the inflammatory response and macrophage activation and their underlying molecular mechanisms are still largely unknown. Moreover, the functional relationship between Syk and MyD88 in the macrophage-activated inflammatory response is poorly understood. Therefore, this study aimed to investigate how Syk and MyD88 cooperate to stimulate the inflammatory response and macrophage function as well as the underlying molecular mechanisms of these biological processes. Here, we show that Syk is a key inducer of reactive oxygen species (ROS)/reactive nitrogen species (RNS) generation by controlling the expression of ROS/RNS-generating enzymes and phagocytosis by suppressing the negative regulator of phagocytosis, suppressor of cytokine signaling 1 (SOCS1), in the macrophage-mediated inflammatory response. These biological functions of Syk are facilitated by MyD88-mediated Syk activation via their interaction, and MyD88 is activated by Src-mediated phosphorylation at the tyrosine 58 residue. Moreover, Src is activated by Ras-related C3 botulinum toxin substrate 1 (Rac1) and polymeric filamentous actin (F-actin).

## Results

### Syk was essential for ROS/RNS generation and consequent phagocytic activity in macrophages

The role of Syk in ROS/RNS generation was investigated in RAW264.7 cells. Since LPS induces ROS/RNS generation in macrophages (de Souza et al, 2007), RAW264.7 cells were treated with LPS and ROS/RNS generation was examined in the wild type (WT) and Syk-deficient RAW264.7 cells. ROS/RNS generation was time-dependently increased by LPS up to 25 to 35 min and decreased after 35 min (Fig. S1A). To examine the role of Syk on ROS/RNS generation in macrophages, Syk knock-out (Syk^-/-^) RAW264.7 cells were generated (Fig. S1B) and LPS-induced ROS/RNS generation was compared with that in the WT RAW264.7 cells.

LPS-induced ROS/RNS generation was significantly inhibited in the Syk^-/-^ RAW264.7 cells compared to the control WT cells at 10, 20, and 30 min (Fig. 1A). LPS-induced ROS generation was also significantly inhibited in Syk siRNA (siSyk)-transfected RAW264.7 cells (Fig. 1B and Fig. S1C) and RAW264.7 cells treated with piceatannol (Pic), a selective Syk inhibitor (Park et al, 2017; Seow et al, 2002), at 10, 20, and 30 min (Fig. 1C).

**Figure 1.**
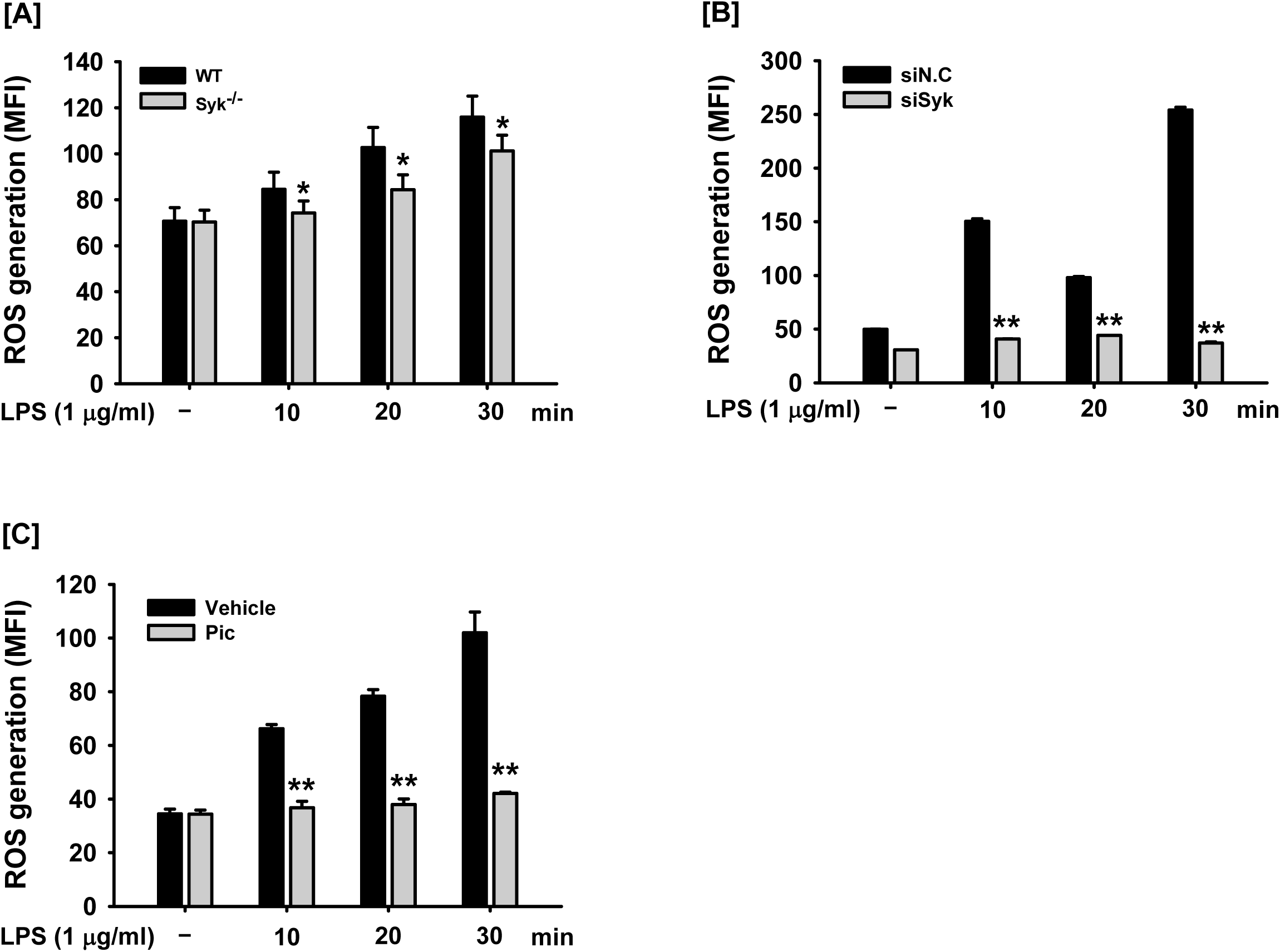
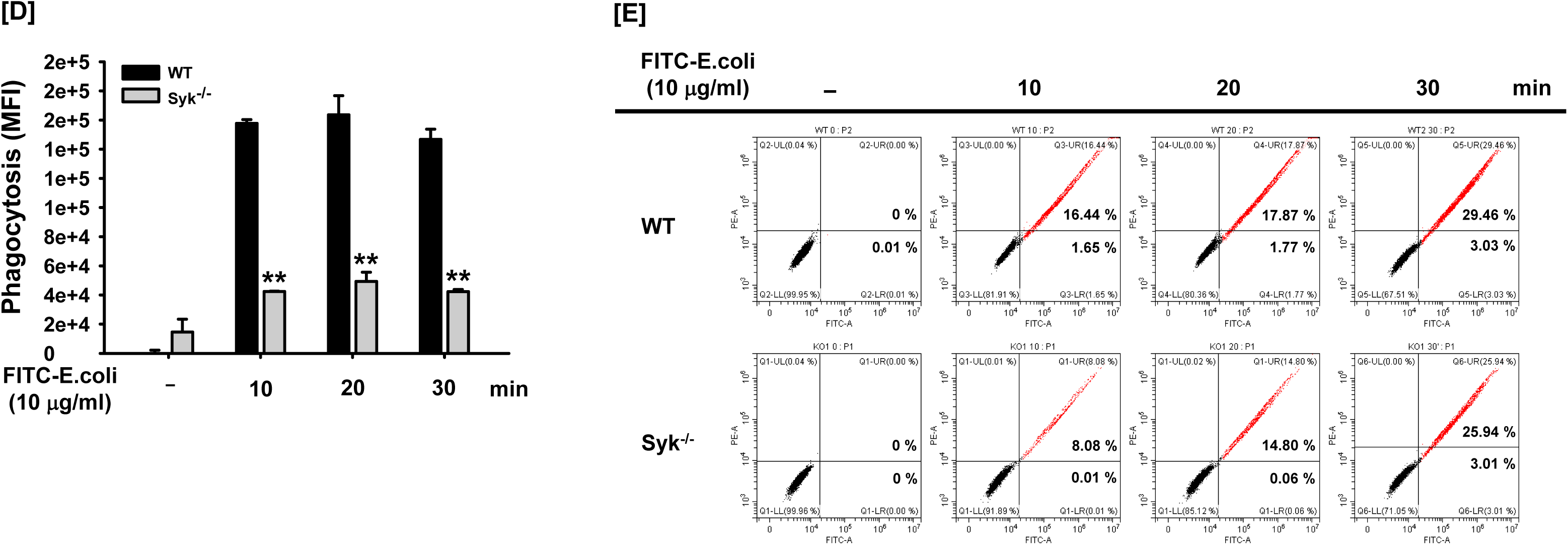
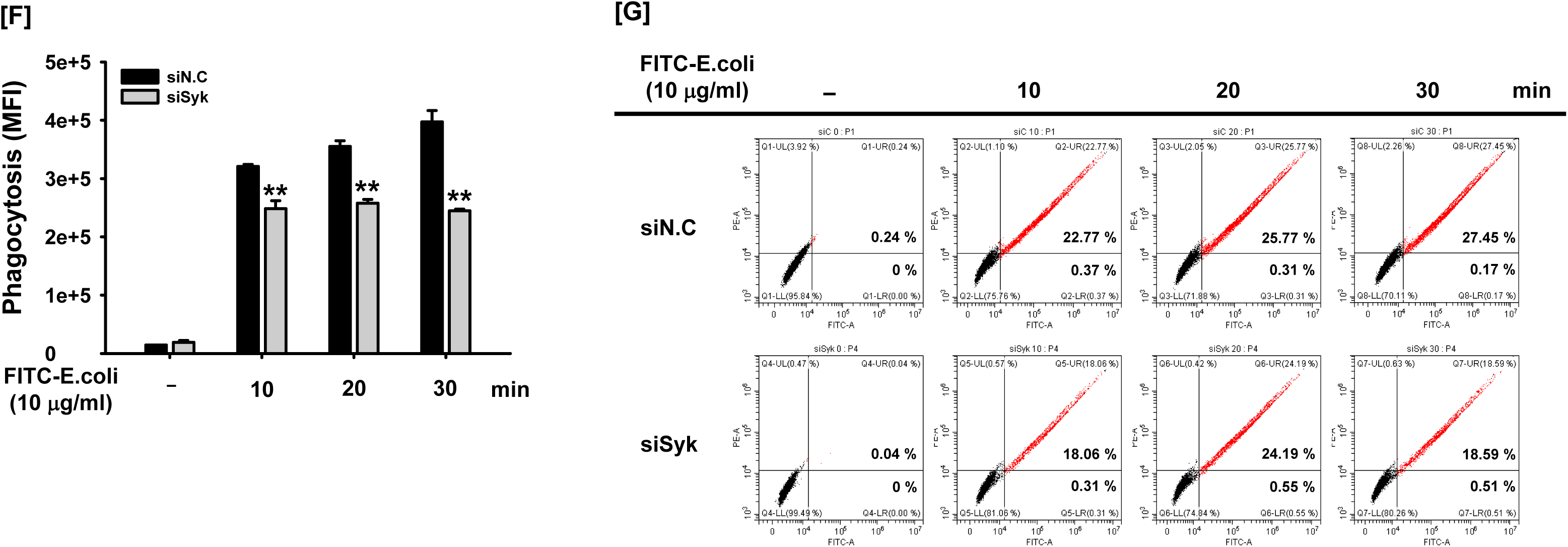
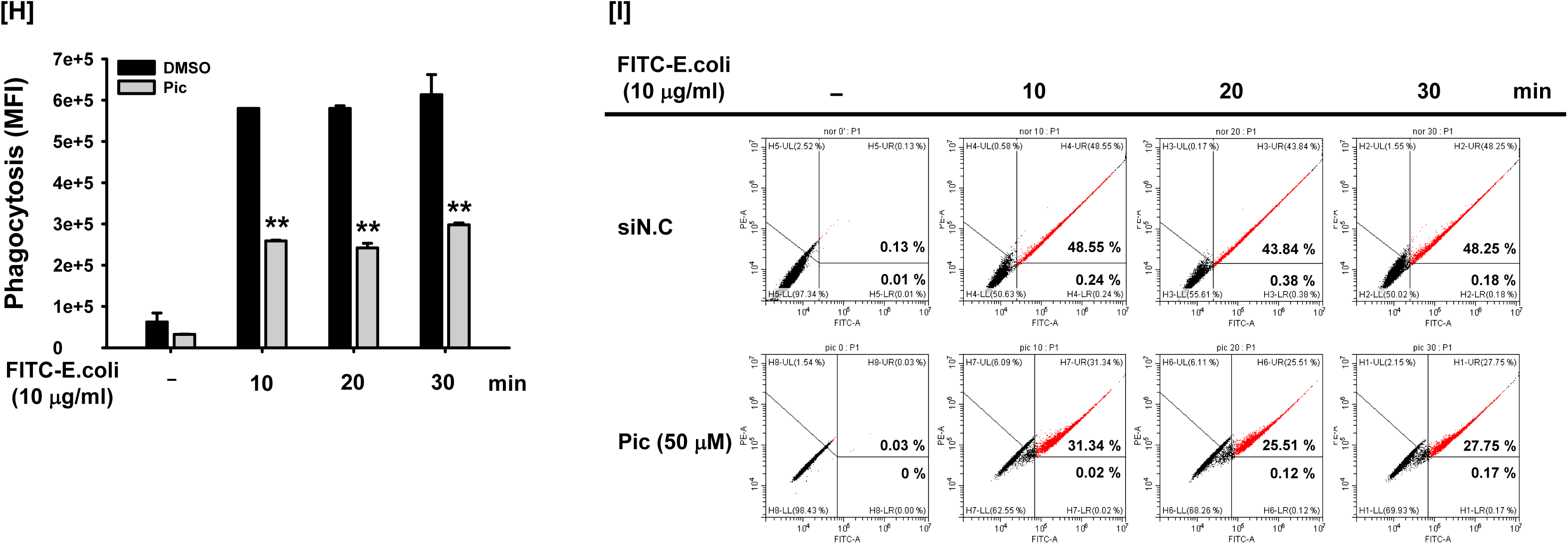
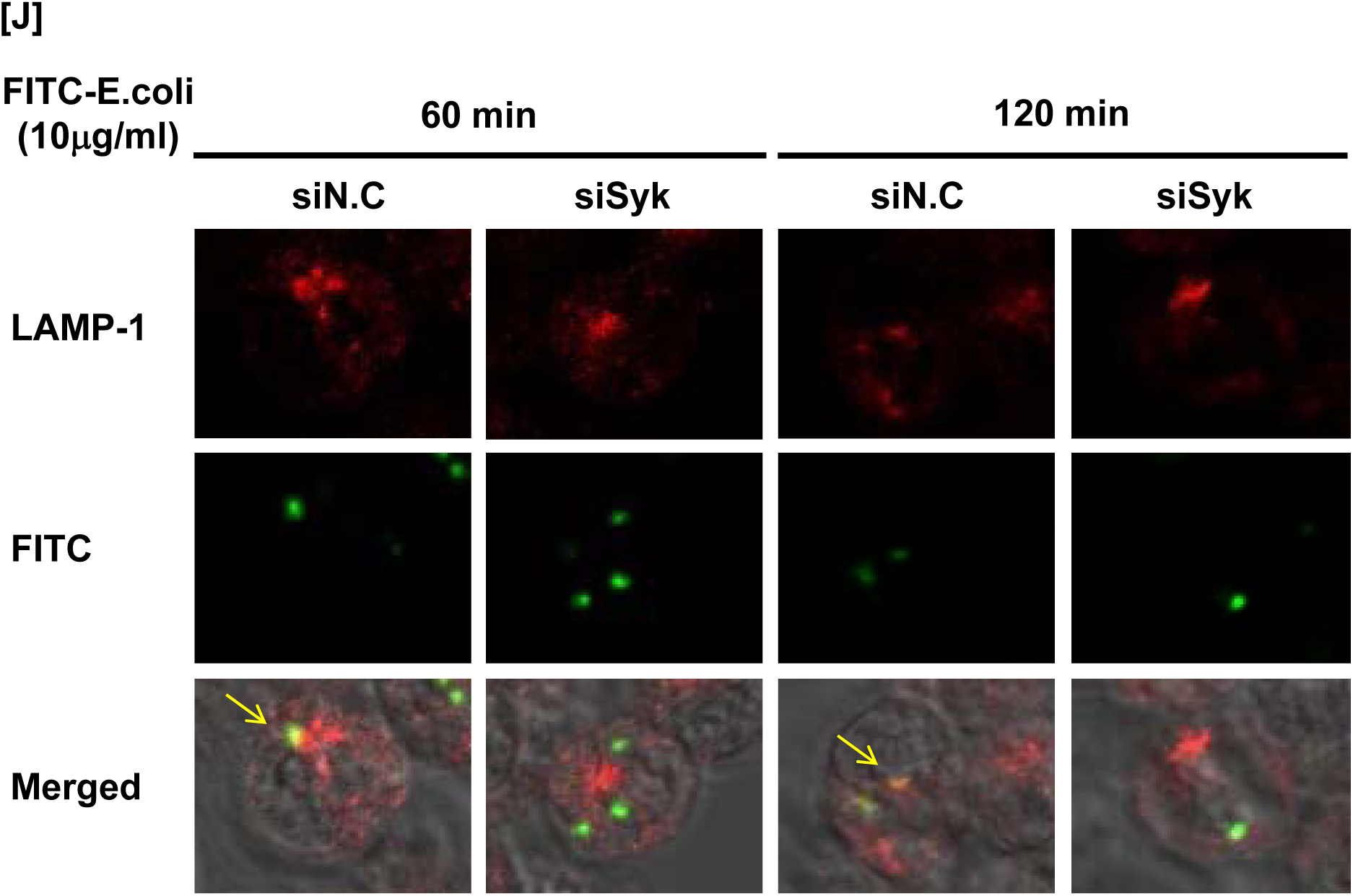
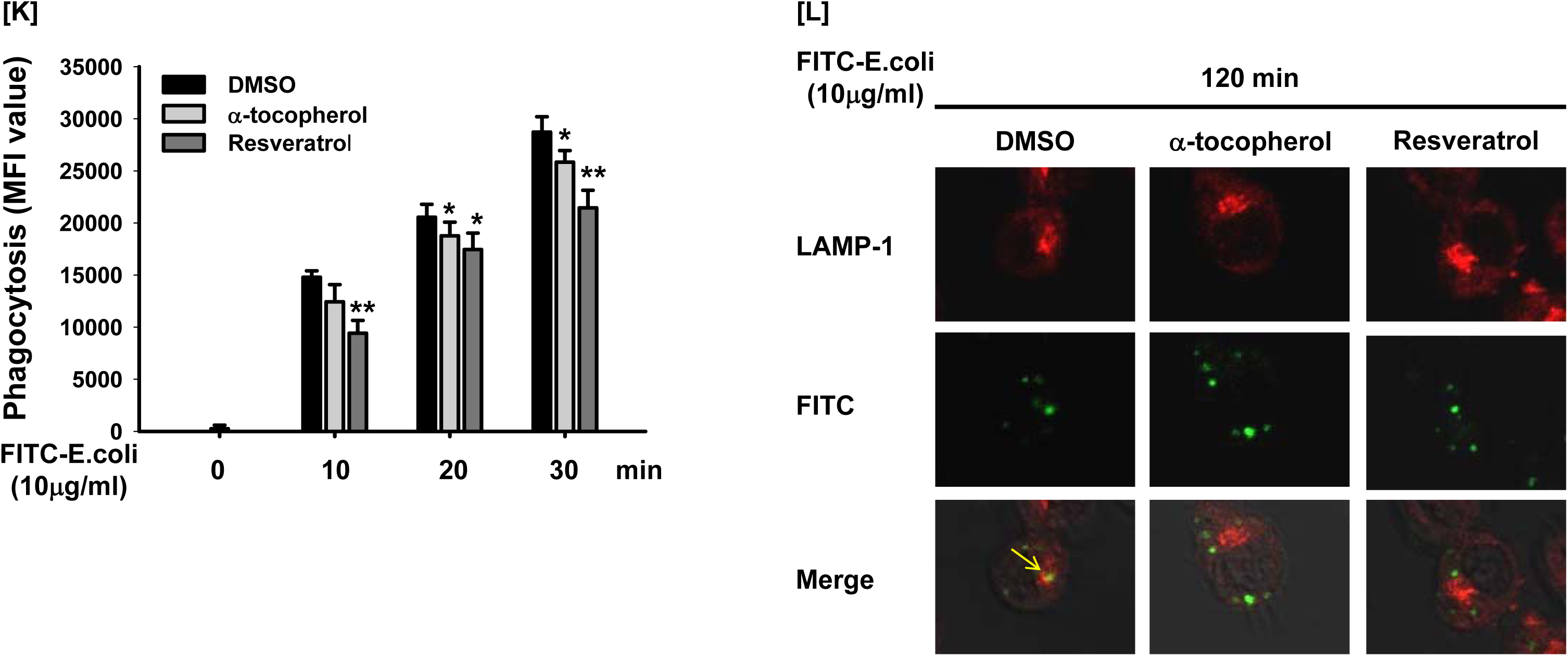
Syk was essential for ROS generation and consequent phagocytic activity in macrophages. (A) WT and Syk^-/-^ RAW264.7 cells, (B) siN.C- and siSyk-transfected RAW264.7 cells, and (C) vehicle (DMSO)- and Pic (50 μM)-treated RAW264.7 cells treated with LPS (1 μg/ml) for the indicated time were incubated with DHR 123 (20 μM) for 20 min, and the ROS level was determined by measuring fluorescence. WT RAW264.7 and Syk^-/-^ RAW264.7 cells were incubated with FITC-E.coli (10 μg/ml) for the indicated time, and the fluorescence was determined by (D) a fluorescence plate reader and (E) a flow cytometer. siN.C-transfected and siSyk-transfected RAW264.7 cells were incubated with FITC-E.coli (10 μg/ml) for the indicated time, and the fluorescence was determined by (F) a fluorescence plate reader and (G) a flow cytometer. Vehicle (DMSO)-treated and Pic-treated RAW264.7 cells were incubated with FITC-E.coli (10 μg/ml) for the indicated time, and the fluorescence was determined by (H) a fluorescence plate reader and (I) a flow cytometer. (J) siN.C-transfected and siSyk-transfected RAW264.7 cells were incubated with FITC-E.coli (10 μg/ml) for the indicated time, and the fluorescence of LAMP-1 and FITC was visualized by confocal microscopy. (K) RAW264.7 cells treated with vehicle (DMSO), α-tocopherol (100 mM), and resveratrol (50 mM) for 12 h were incubated with FITC-E.coli (10 μg/ml) for the indicated time, and the fluorescence was determined by a fluorescence plate reader. (L) RAW264.7 cells treated with vehicle (DMSO), α-tocopherol (100 mM), and resveratrol (50 mM) for 12 h were incubated with FITC-E.coli (10 μg/ml) for 120 min, and the fluorescence of LAMP-1 and FITC was visualized by confocal microscopy. Arrow indicates the overlap of LAMP-1 and FITC. N.C.: Negative control. \**P* < 0.05, \*\**P* < 0.01 compared to controls.

Next, the role of Syk on the phagocytic activity of macrophages was investigated. Phagocytic activity was time-dependently increased in the WT RAW264.7 cells, while it was markedly suppressed in the Syk^-/-^ RAW264.7 cells compared to WT cells at 10, 20, and 30 min (Fig. 1D and E). Phagocytic activity was also markedly suppressed in the siSyk-transfected RAW264.7 cells (Fig. 1F and G) and the RAW264.7 cells treated with Pic at 10, 20, and 30 min (Fig. 1H and I). Confocal microscopic analysis further confirmed that phagocytic activity was decreased in the siSyk-transfected RAW264.7 cells compared to control WT cells at 60 and 120 min, and the phagocytosis occurred in a lysosome-dependent manner (Fig. 1J).

To examine whether ROS/RNS generation induces the phagocytic activity of macrophage, RAW264.7 cells were treated with either α-tocopherol or resveratrol, both of which are known to inhibit ROS/RNS generation (Cheng et al, 2016; Lin et al, 2018; Thews et al, 2005; Wu et al, 2014), and phagocytic activity of these cells was determined. Phagocytic activity was inhibited by both α-tocopherol and resveratrol in RAW264.7 cells at 10 to 30 min (Fig. 1K) as well as 120 min in a lysosome-dependent manner (Fig. 1L). These results strongly suggest that Syk rapidly (within 30 min) induces ROS//RNS generation, which in turn increases the phagocytic activity of macrophages.

### Syk induced the phagocytosis activity through expression of ROS/RNS-generating enzymes and tyrosine nitration of SOCS1

The molecular mechanism of Syk-induced ROS/RNS generation in macrophages was investigated. First, the role of Syk on the gene expression of ROS-generating enzymes in the LPS-stimulated RAW264.7 cells was examined. mRNA expression of the ROS/RNS-generating enzyme iNOS, the inflammatory enzyme COX-2, and the pro-inflammatory cytokines TNF-α, IL-1β, and IL-6 was upregulated by LPS in RAW264.7 cells (Fig. S2). Since ROS/RNS generation gradually increased and was highest at 25 to 30 min after LPS treatment (Fig. S1A), mRNA expression of ROS/RNS-generating enzymes was examined in the RAW264.7 cells treated with LPS for up to 30 min. mRNA expression of the ROS/RNS-generating enzymes including iNOS, NADPH oxidase 1 (NOX1), aldehyde oxidase 1 (AOX1), and xanthine dehydrogenase (XDH) was highly increased by LPS at 10 min and decreased at 20 and 30 min, whereas mRNA expression of amine oxidase copper containing-3 (AOC3) was increased by LPS only at 30 min in RAW264.7 cells (Fig. 2A). However, LPS-induced mRNA expression of these enzymes was significantly suppressed in the Syk^-/-^ RAW264.7 cells compared to the control WT cells (Fig. 2B). LPS-induced mRNA expression of these enzymes was also suppressed in the siSyk-transfected RAW264.7 cells (Fig. 2C) and the RAW264.7 cells treated with Pic (Fig. 2D). Then, we studied how ROS/RNS produced by these enzymes regulate phagocytosis in macrophages. Interestingly, tyrosine-nitrated (N-Tyr) proteins with molecular weights around 98 kDa, 39 kDa, and 23 kDa were identified in the LPS-stimulated RAW264.7 cells from protein nitration profile screening, and their N-Tyr levels were gradually increased, with the highest levels at 30 to 40 min (Fig. S3A). Immunoprecipitation and Western blot analysis identified the 23 kDa N-Tyr protein in the LPS-stimulated RAW264.7 cells as suppressor of cytokine signaling 1 (SOCS1) (Fig. S3B). However, the other two N-Tyr proteins were not identified by immunoprecipitation and Western blot analysis (data not shown). The N-Tyr of SOCS1 was increased by sodium nitroprusside (SNP), a RNS generating reagent, in the SOCS1-transfected RAW264.7 cells (Fig. 2E). On the other hand, the N-Tyr of SOCS1 induced by LPS was clearly eliminated by the iNOS inhibitor or radical scavengers, N(G)-Nitro-L-arginine methyl ester (L-NAME) and α-tocopherol (Jerkic et al, 2012; Thews et al, 2005; Wu et al, 2014) (Fig. 2F). These results suggest that N-Tyr of SOCS is regulated by ROS/RNS in LPS-stimulated macrophages. Since Syk induced phagocytic activity by increasing ROS/RNS generation in RAW264.7 cells (Fig. 1), the role of SOCS1 on the *E.coli*-induced phagocytic activity in macrophages was further examined. Phagocytic activity was significantly reduced in the SOCS1-overexpressed RAW264.7 cells compared to the control WT cells (Fig. 2G). On the contrary, inhibition of SOCS1 expression by SOCS1-specific small interference RNA (siSOCS1) induced the phagocytic activity of RAW264.7 cells (Fig. 2H), indicating that SOCS1 acts as a negative regulator in phagocytosis. Moreover, the phagocytic activity was decreased by SOCS1 in RAW264.7 cells, however, SNP recovered the phagocytic activity decreased by SOCS to that of WT RAW264.7 cells (Fig. 2I). Collectively, ROS/RNS generated under the LPS or *E.coli* infection conditions seems to SOCS1 inactivation through nitration at tyrosine residues, thereby increase phagocytosis.

**Figure 2.**
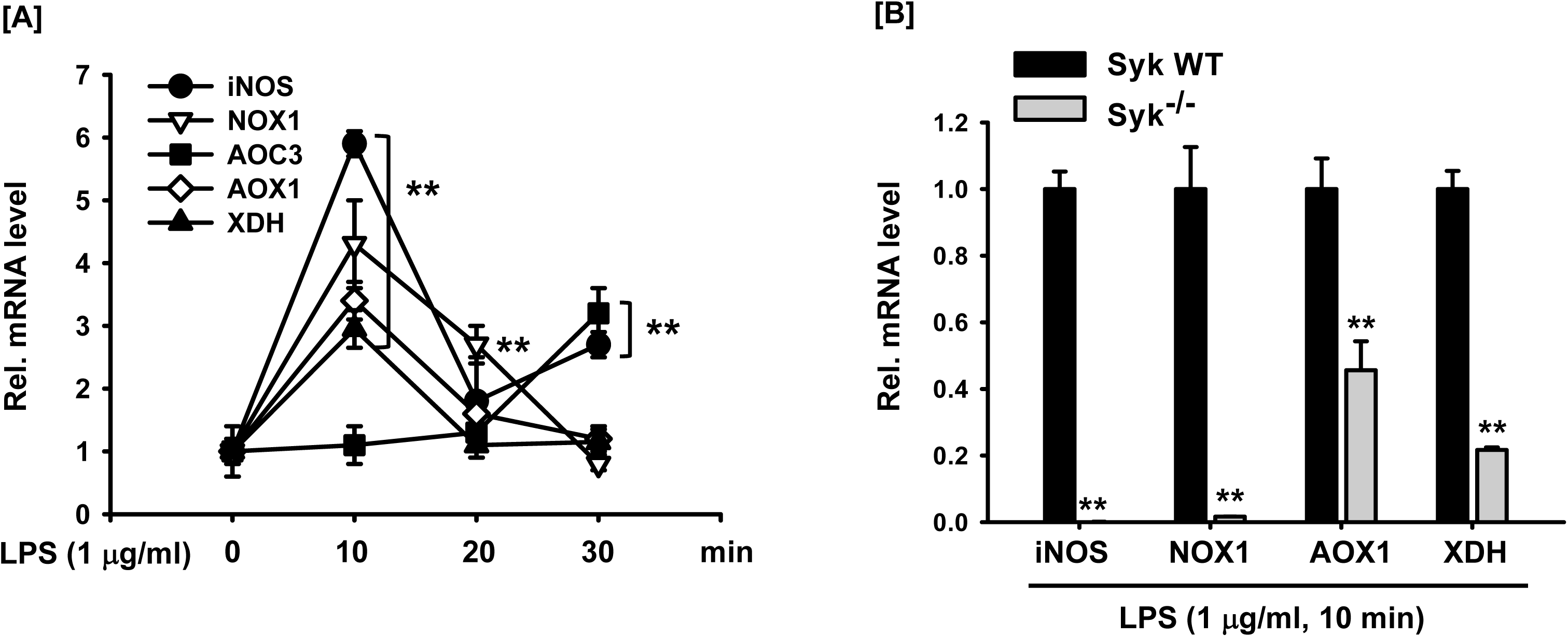
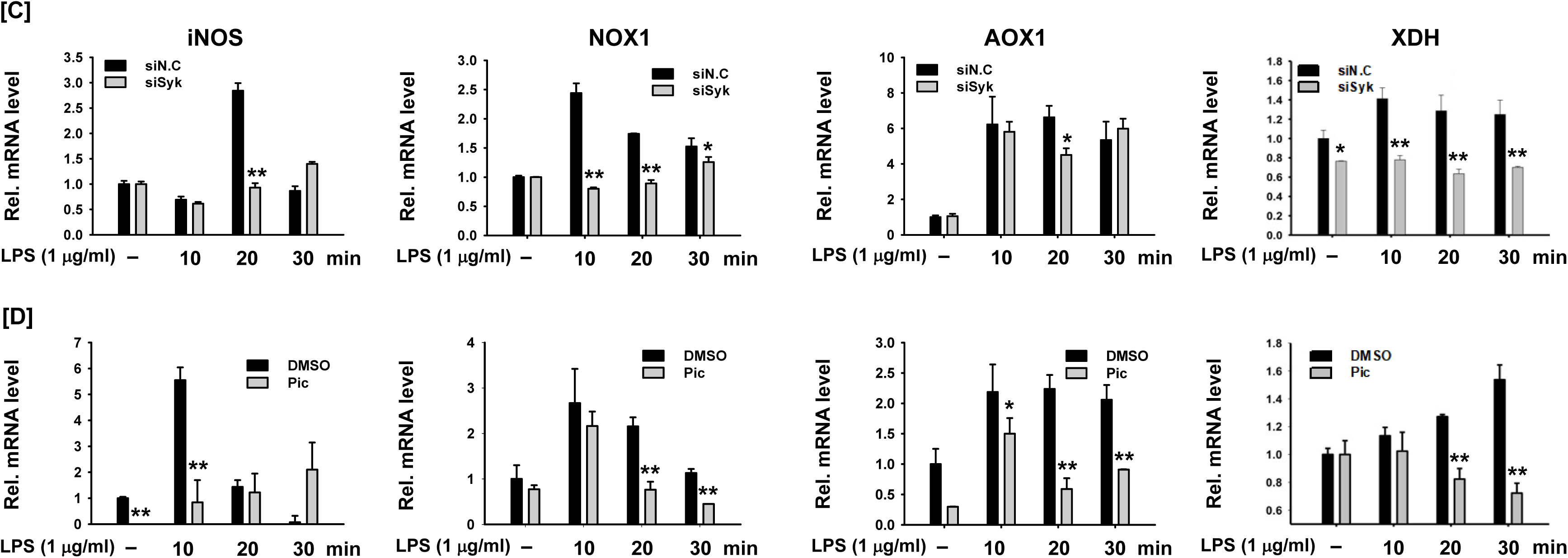
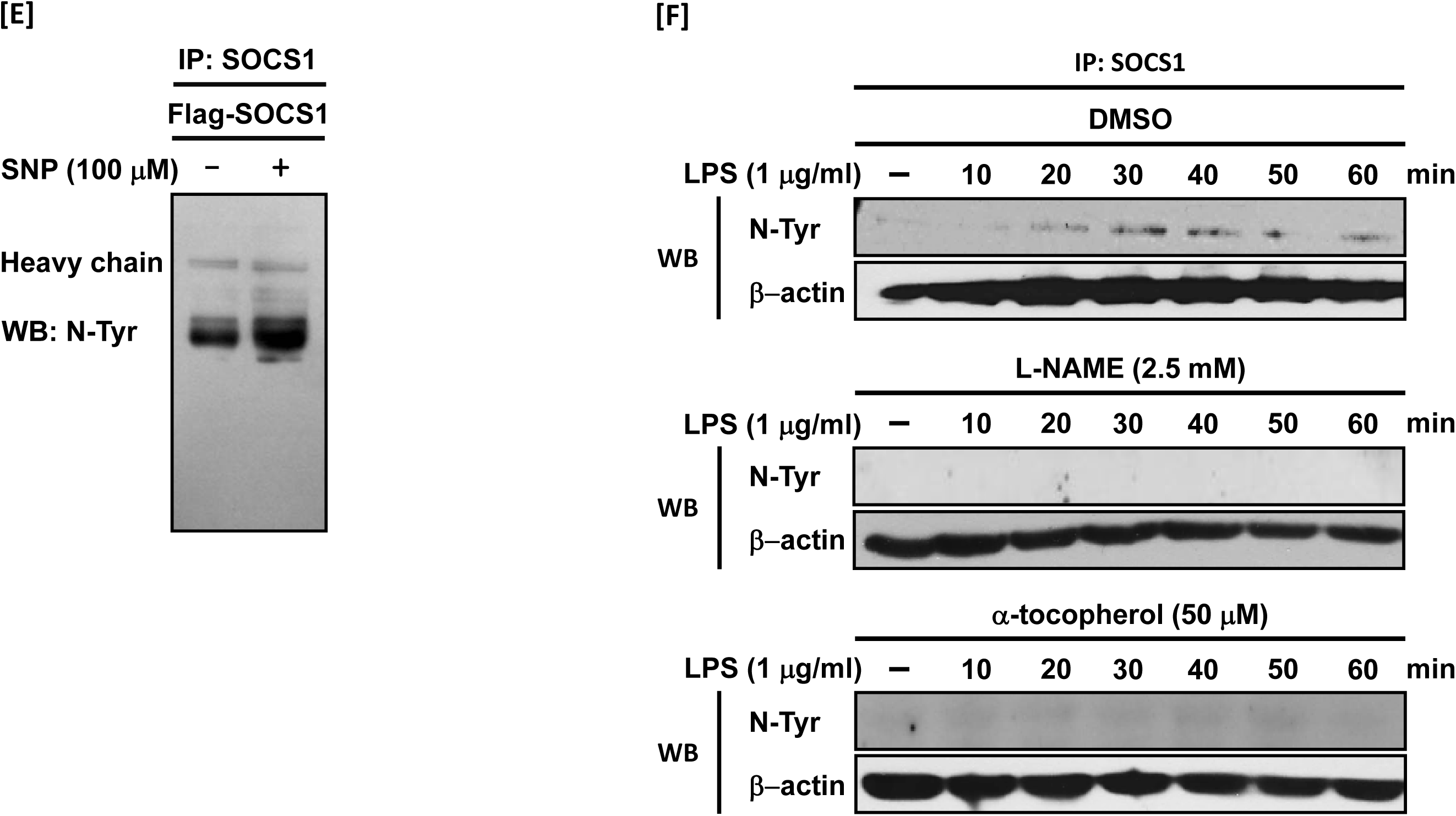
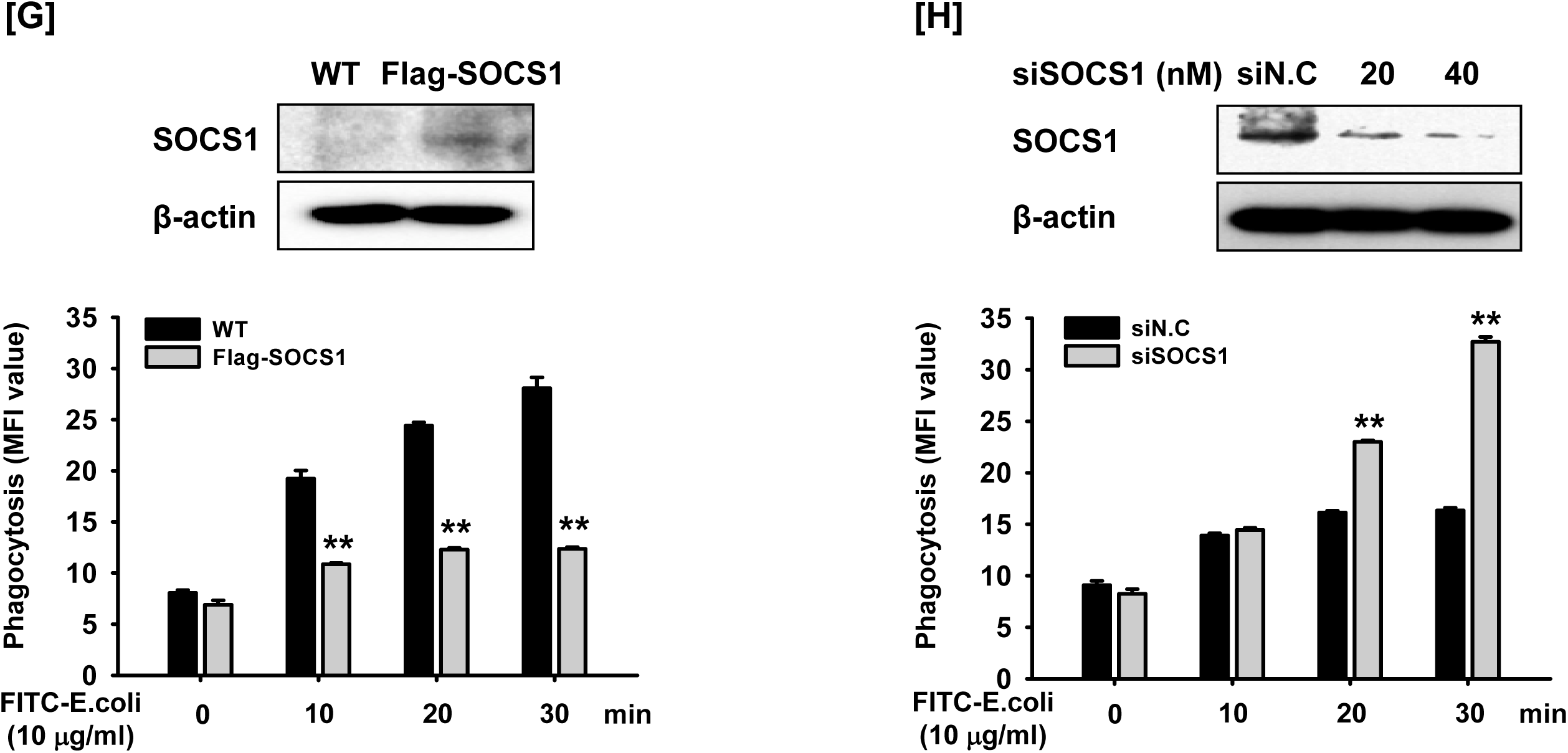
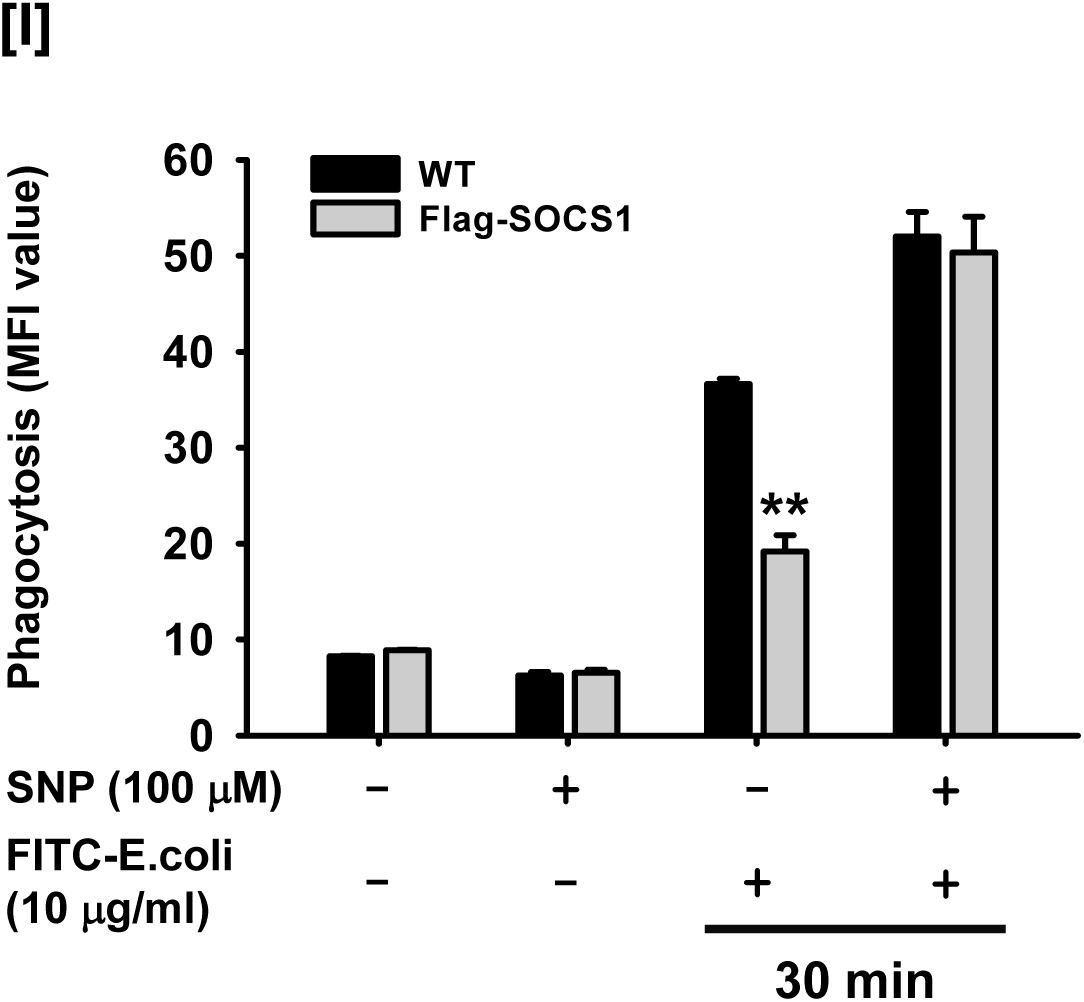
Syk induced the phagocytosis activity through expression of ROS-generating enzymes and tyrosine nitration of SOCS1. (A) RAW264.7 cells were treated with LPS (1 μg/ml) for the indicated time, and mRNA expression of the target genes was determined by quantitative real time PCR. (B) WT RAW264.7 and Syk^-/-^ RAW264.7 cells were treated with LPS (1 μg/ml) for 10 min, and mRNA expression of the target genes was determined by quantitative real time PCR. (C) siN.C-transfected and siSyk-transfected RAW264.7 cells and (D) vehicle (DMSO) and Pic-treated RAW 264.7 cells were treated with LPS (1 μg/ml) for the indicated time, and mRNA expression of the target genes was determined by quantitative real time PCR. (E) HEK293 cells transfected with SOCS1 plasmid for 48 h were treated with either vehicle (DMSO) or SNP (100 μM) for 3 h, and N-Tyr SOCS1 was detected by immunoprecipitation and Western blot analysis. (F) N-Tyr SOCS1 in the whole cell lysates of RAW264.7 cells treated with either vehicle (DMSO, 12 h), L-NAME (2.5 mM, 1 h), and α-tocopherol (100 mM, 12 h) for 12 h, followed by treatment of LPS (1 μg/ml) for the indicated time was detected by immunoprecipitation and Western blot analysis. (G) RAW264.7 cells were transfected with the SOCS1-expression plasmid 48 h, and SOCS1 expression was detected by Western blot analysis. RAW264.7 cells transfected with either empty (pcDNA) or SOCS1 plasmid for 48 h were incubated with FITC-E.coli (10 μg/ml) for the indicated time, and the fluorescence was determined by a fluorescence plate reader. (H) RAW264.7 cells were transfected with siN.C or siSOCS1 (20, 40 nM) for 24 h, and SOCS1 expression was detected by Western blot analysis. RAW264.7 cells transfected with siN.C or siSOCS1 (20 nM) for 24 h were incubated with FITC-E.coli (10 μg/ml) for the indicated time, and the fluorescence was determined by a fluorescence plate reader. (I) RAW264.7 cells treated with SNP (100 μM) for 3 h were incubated with FITC-E.coli (10 μg/ml) for 30 min, and the fluorescence was determined by a fluorescence plate reader. \**P* < 0.05, \*\**P* < 0.01 compared to controls.

### Syk induced the NF-κB signaling pathway via MyD88

Next, the underlying molecular mechanism associated with Syk signaling in LPS-stimulated macrophages was investigated. Syk was immediately phosphorylated by LPS at 1 min, and the phosphorylation of p85 and IκBα followed with the highest levels at 3 min and 5 min, respectively (Fig. S4A). IκBα was markedly phosphorylated by LPS at 5 min and then degraded from 15 min to 30 min (Fig. S4B). In accordance with IκBα degradation, nuclear translocation of the NF-κB subunits p65 and p50 was initiated after 5 min by LPS and was highest at 15 min lasting to 30 min (Fig. S4C). IκBα phosphorylation was notably delayed in the Syk^-/-^ RAW264.7 cells compared to the control WT cells (Fig. 3A), and the nuclear translocation of p65 and p50 was also markedly delayed in the Syk^-/-^ RAW264.7 cells compared to the control WT cells (Fig. 3B). These results were confirmed by the inhibition of IκBα phosphorylation and nuclear translocation of p65 and p50 in the RAW264.7 cells treated with Pic (Fig. 3C and D). Syk more strongly induced the NF-κB-mediated luciferase reporter gene activity than Src and Akt1 in the RAW264.7 cells under the NF-κB stimulators LPS, TNF-α, and PMA (Fig. 3E). The NF-κB-mediated luciferase reporter gene activity induced by LPS was significantly suppressed in the Syk^-/-^ RAW264.7 cells compared to the control WT cells (Fig. 3F), and Syk inhibition by its selective inhibitor Pic significantly reduced the NF-κB-mediated luciferase reporter gene activity in RAW264.7 cells (Fig. S4D). These results indicate that Syk is a key regulator of NF-κB signaling in LPS-treated conditions.

**Figure 3.**
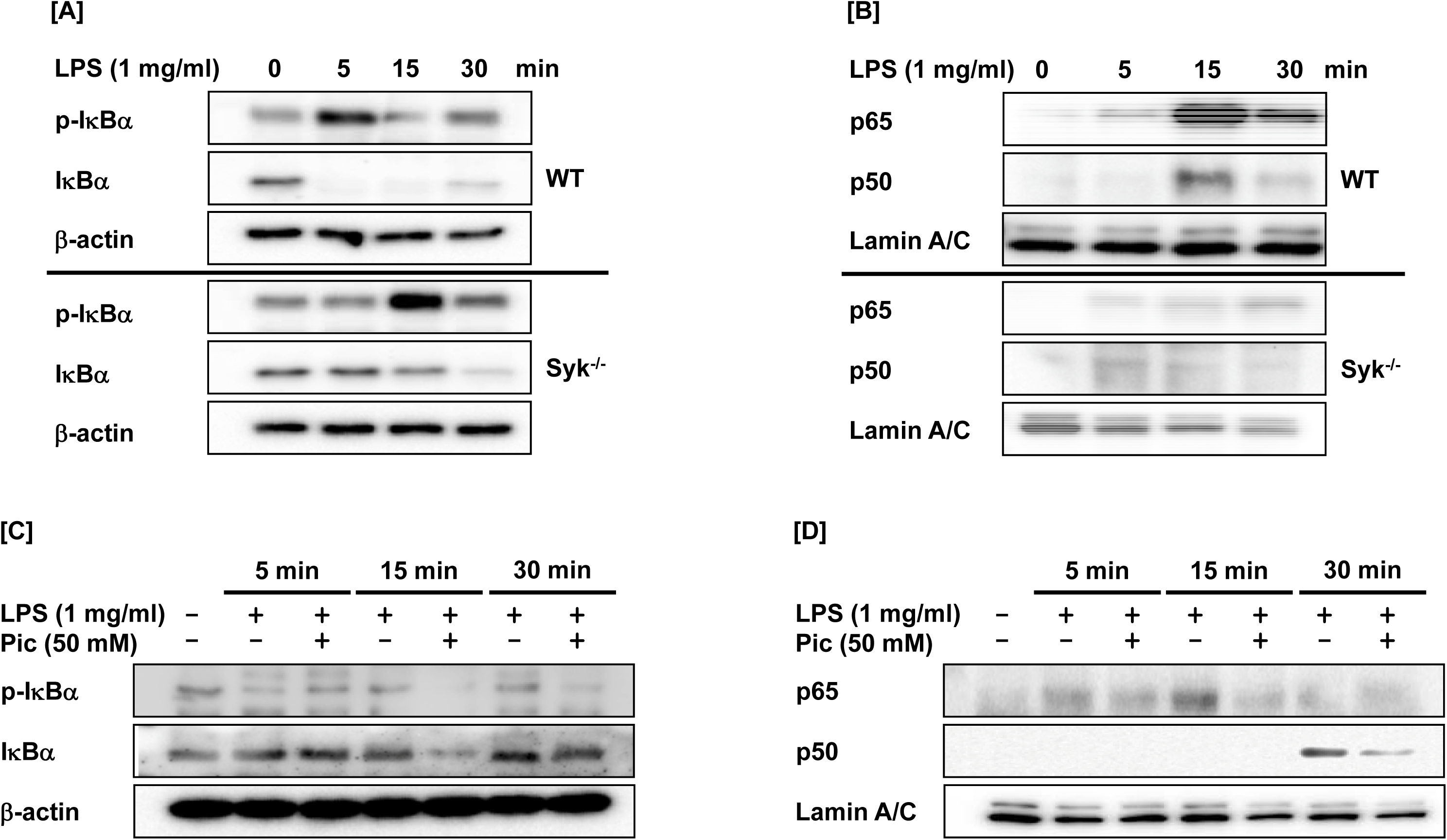
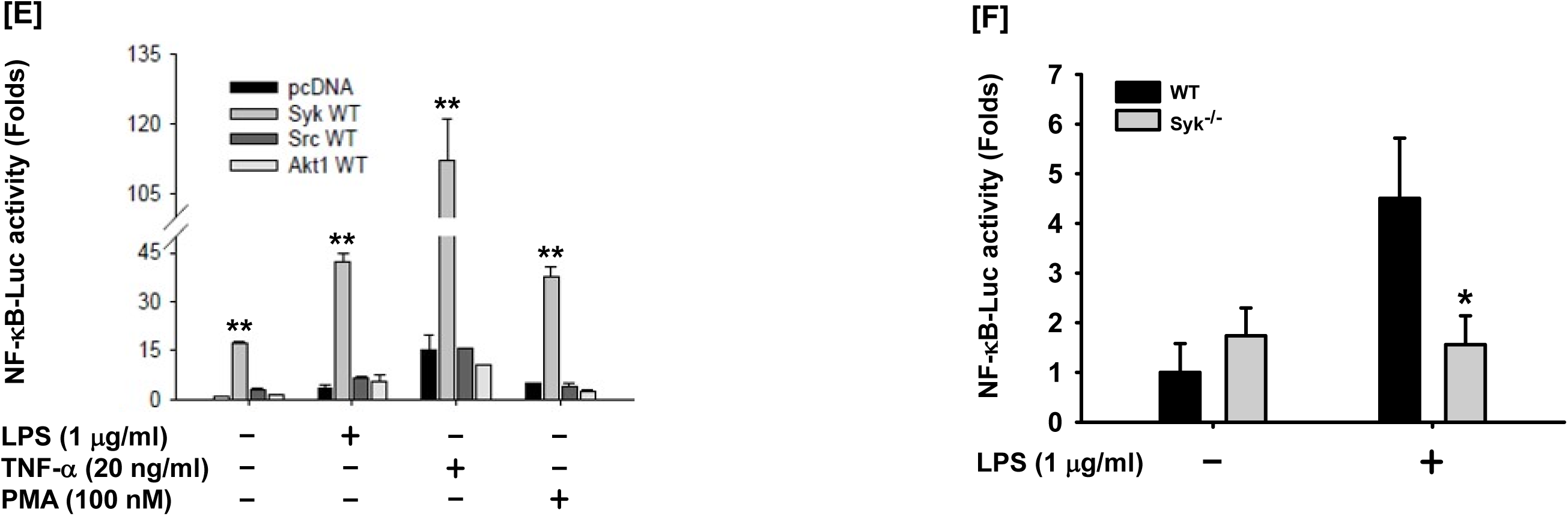
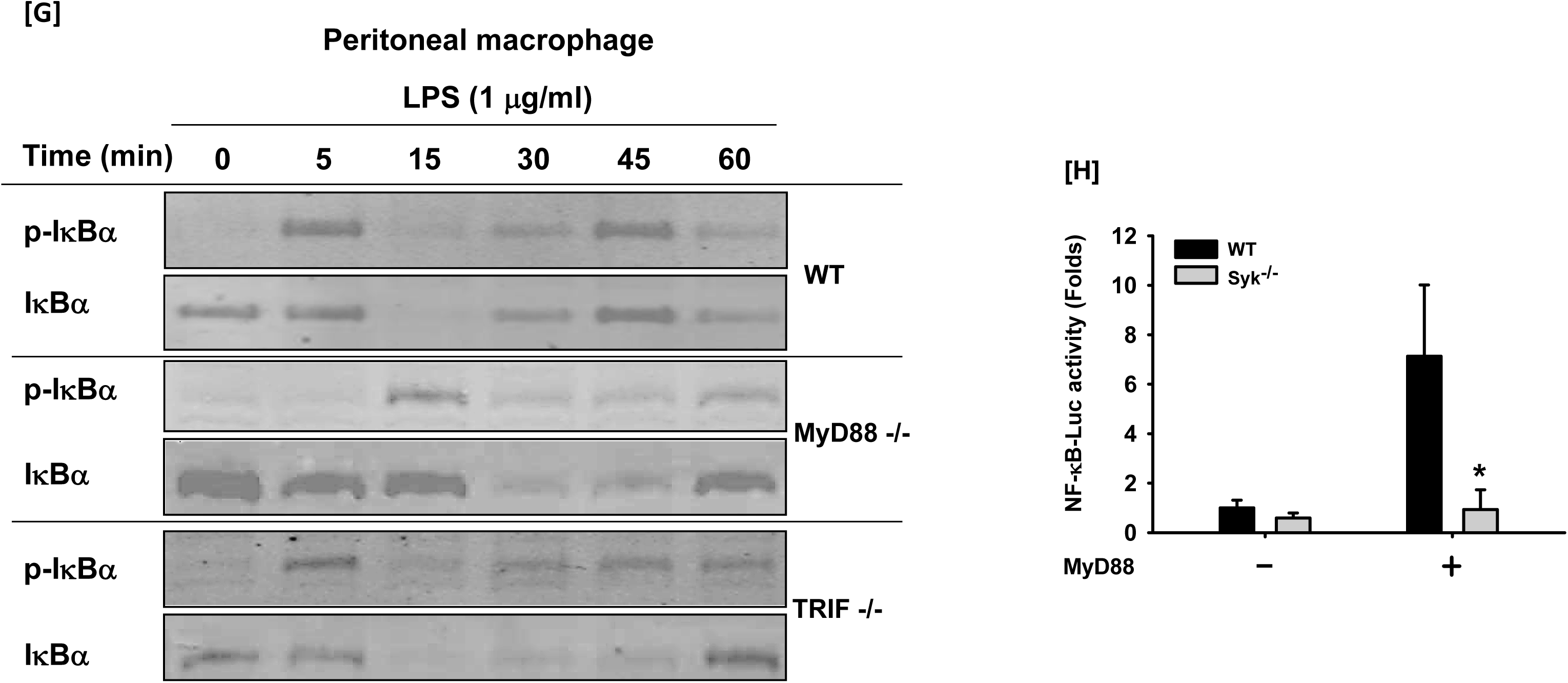
Syk induced the NF-κB signaling pathway via MyD88. (A) IκBα and p-IκBα in the whole cell lysates of WT RAW264.7 and Syk^-/-^ RAW264.7 cells treated with LPS (1 μg/ml) for the indicated time were detected by Western blot analysis. (B) p65 and p50 in the nuclear lysates of WT RAW264.7 and Syk^-/-^ RAW264.7 cells treated with LPS (1 μg/ml) for the indicated time were detected by Western blot analysis. (C) IκBα and p-IκBα in the whole cell lysates of RAW264.7 cells treated with LPS (1 μg/ml) for the indicated time in the absence or presence of Pic (50 mM) were determined by Western blot analysis. (D) p65 and p50 in the nuclear lysates of RAW264.7 cells treated with LPS (1 μg/ml) for the indicated time in the absence or presence of Pic (50 mM) were determined by Western blot analysis. (E) RAW264.7 cells co-transfected with the NF-κB-Luciferase reporter gene plasmid and either empty (pcDNA), Syk, Src, or Akt plasmid were treated with either LPS (1 μg/ml), TNF-α (20 ng/ml), or PMA (100 nM) for 24 h, and the luciferase activity was measured by a luminometer and normalized to β-galactosidase activity. (F) WT RAW264.7 and Syk^-/-^ RAW264.7 cells transfected with the NF-κB-Luciferase reporter gene plasmid for 24 h were treated with LPS (1 μg/ml) for 24 h. (G) ΙκBα, and p-IκBα were detected by Western blot analysis in the peritoneal macrophages extracted from WT, MyD88-/-, and TRIF-/- mice under LPS (1 μg/ml) treatment condition. (H) WT RAW264.7 and Syk^-/-^ RAW264.7 cells were co-transfected with the NF-κB-Luciferase reporter gene plasmid and MyD88 plasmid for 48 h. The luciferase activity of these cells was measured by a luminometer and normalized to β-galactosidase activity. \**P* < 0.05, \*\**P* < 0.01 compared to controls.

MyD88 and TRIF are the adaptors that bind to the cytoplasmic domains of TLR4, the molecular receptor of LPS, and induce the activation of TLR4-mediated NF-κB signaling pathways during the innate immune response (Takeda & Akira, 2005; Yi et al, 2014). Interestingly, LPS-induced IκBα phosphorylation was markedly delayed only in the MyD88^-/-^ peritoneal macrophages, not TRIF-/- cells (Fig. 3G). Moreover, the phosphorylation of IκBα at 5 min was observed under treatments of Pam3CSK4 stimulating TLR1 and TLR2, and LPS, a stimulator of TLR4 (Fig. S4E). In here, TLR1 and 2 have only the MyD88 as an adapter molecule. On the other hand, Poly I:C, activating TLR3 signaling via TRIF, induced the phosphorylation of IκBα at 60 min (Fig. S4E). NF-κB luciferase activity induced by MyD88 was also inhibited in the Syk^-/-^ RAW 264.7 cells compared to WT cells (Fig. 3H). These results suggest that Syk, which is activated at the initial time under the LPS-stimulation, is closely involved in MyD88-dependent NF-κB signals. Additionally, MyD88^-/-^ RAW 264.7 cells showed inhibition of ROS/RNS generation, phagocytosis, and expression of ROS/RNS-producing enzymes, as observed in Syk^-/-^cells, indicating that MyD88 and Syk signals are closely related to each other (Fig. S4F-I).

### MyD88 interacted with and activated Syk

The mechanism by which Syk is activated in macrophages was investigated. Since MyD88 is an upstream molecule of Syk in the TLR4-mediated inflammatory responses (Fig. 3G and H), and ROS/RNS generation, phagocytic activity, and expression of ROS/RNS-generating enzymes were significantly reduced in the MyD88^-/-^ RAW264.7 cells (Fig. S4F – I), the relationship between MyD88 and Syk was examined. Co-transfection of HEK293 cells with Flag-MyD88 and Myc-Syk revealed that MyD88 increased the phosphorylation of Syk (Fig. 4A). In addition, MyD88 markedly induced NF-κB-mediated luciferase reporter gene activity in RAW264.7 cells, while the activity was dramatically reduced in the RAW264.7 cells transfected with the construct of Syk kinase domain deletion mutant (Myc-Syk(ΔKD)) (Fig. 4B).

**Figure 4.**
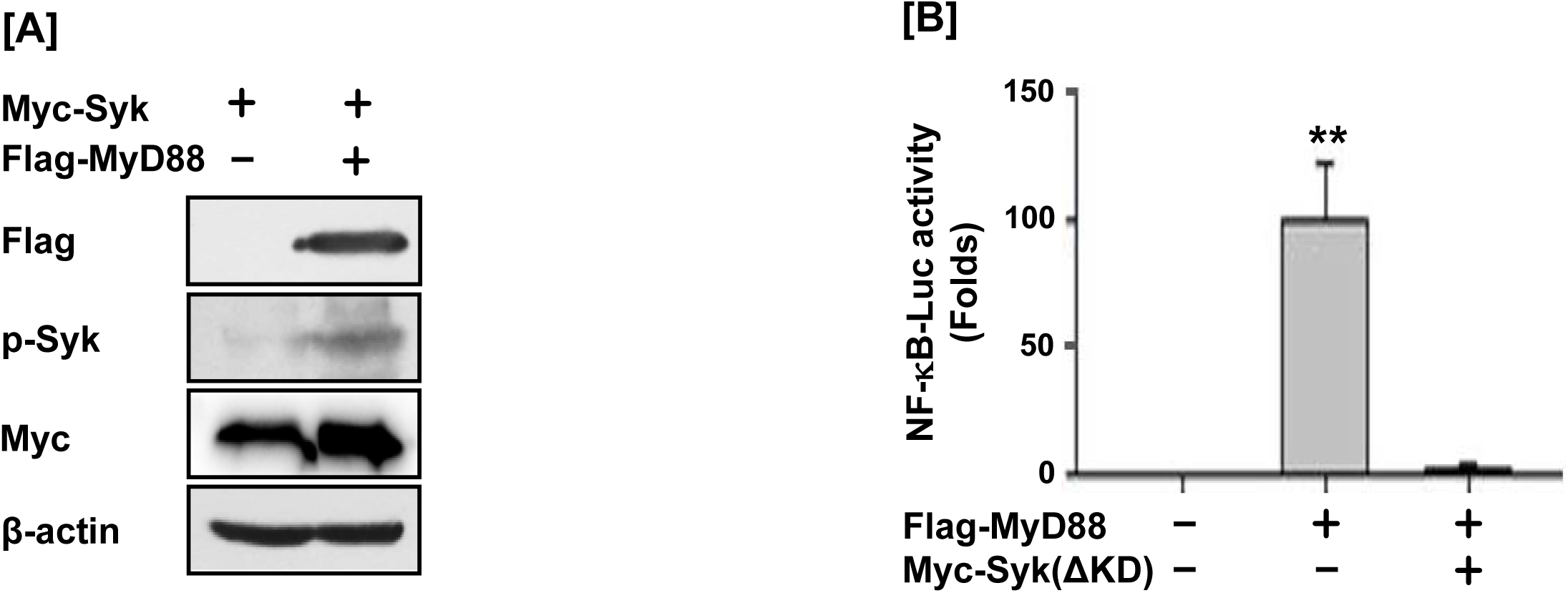
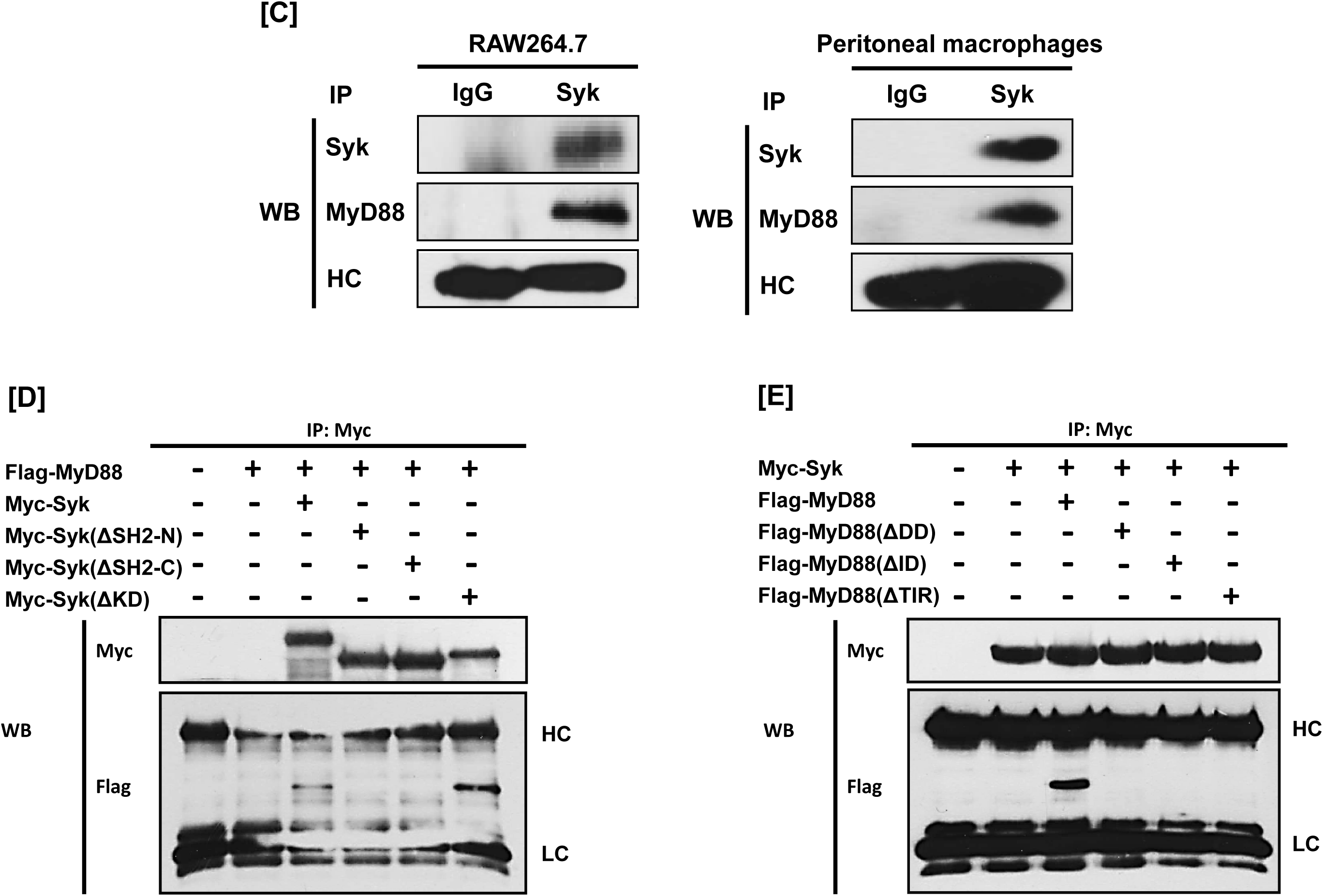
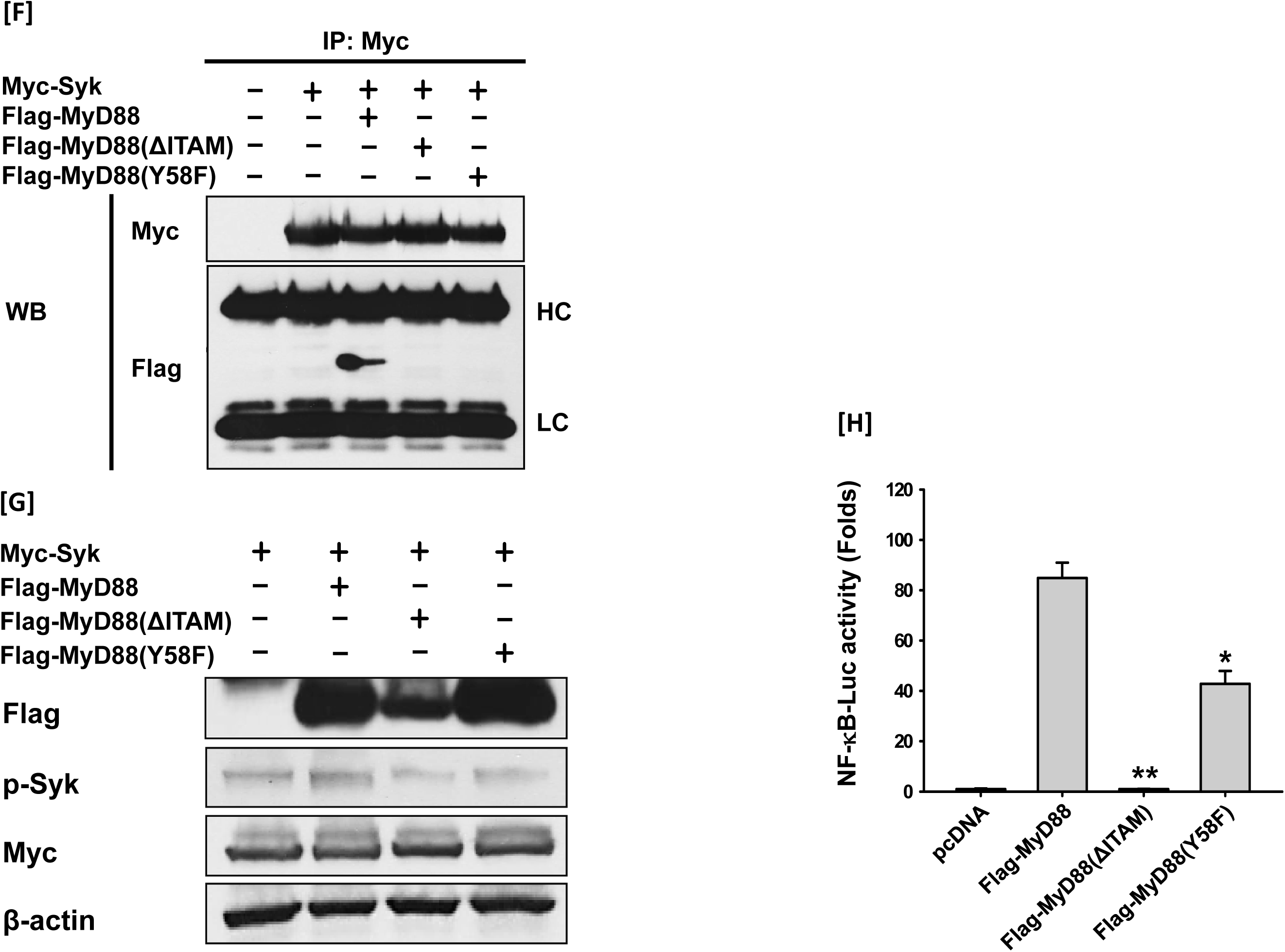
MyD88 interacted with and activated Syk. (A) p-Syk, Flag, and Myc were detected by Western blot analysis in the whole cell lysates of HEK293 cells which were co-transfected with Myc-Syk and either empty (pcDNA) or Flag-MyD88 plasmid for 48 h (B) HEK293 cells co-transfected with the NF-κB-Luciferase reporter gene plasmid, either empty (pcDNA) or Flag-MyD88 plasmids, and empty (pcDNA) or Myc-Syk ΔKD plasmids for 48 h, and the luciferase activity was measured by a luminometer and normalized to β-galactosidase activity. (C) Whole cell lysates of RAW264.7 cells and peritoneal macrophages were immunoprecipitated with either non-specific control immunoglobulin G (IgG) or Syk antibodies, followed by detection of Syk and MyD88 by Western blot analysis. (D) Whole cell lysates of HEK293 cells co-transfected with Flag-MyD88 and either Myc-Syk or Syk deletion mutant plasmids for 48 h were immunoprecipitated with Myc antibodies, followed by detection of Myc and Flag by Western blot analysis. (E) Whole cell lysates of HEK293 cells co-transfected with Myc-Syk and either Flag-MyD88 or MyD88 deletion mutant plasmids for 48 h were immunoprecipitated with Myc antibodies, followed by detection of Myc and Flag by Western blot analysis. (F) Whole cell lysates of HEK293 cells co-transfected with Myc-Syk and either Flag-MyD88 or Flag-MyD88(ΔITAM) and Flag-MyD88(Y58F) plasmids for 48 h were immunoprecipitated with Myc antibodies, followed by detection of Myc and Flag by Western blot analysis. (G) p-Syk, Flag, and Myc in the whole cell lysates of HEK293 cells co-transfected with Myc-Syk and either Flag-MyD88 or Flag-MyD88(ΔITAM) and Flag-MyD88(Y58F) plasmids for 48 h were detected by Western blot analysis. (H) HEK293 cells co-transfected with the NF-κB-Luciferase reporter gene plasmid and either empty (pcDNA), Flag-MyD88, Flag-MyD88(ΔITAM) or Flag-MyD88(Y58F) plasmids for 48 h, and the luciferase activity was measured by a luminometer and normalized to β-galactosidase activity. \**P* < 0.05, \*\**P* < 0.01 compared to controls.

Molecular interaction between Syk and MyD88 was examined next. MyD88 interacted with Syk in RAW264.7 cells and peritoneal macrophages (Fig. 4C). For interaction site mapping, domain deletion mutants of Syk (Fig. S5A) and MyD88 (Fig. S5B) were generated, and their interaction was examined by co-transfecting HEK293 cells with the mutant plasmids. MyD88 interacted with WT Syk and Myc-Syk(ΔKD), but did not interact with Syk with the N-terminal Src homology-2 domain deletion (Myc-Syk(ΔSH2-N)) or the C-terminal Src homology-2 domain deletion (Myc-Syk(ΔSH2-C)) (Fig. 4D). Moreover, Syk interacted with WT MyD88, but did not interact with MyD88 with the death domain deletion (Flag-MyD88(ΔDD)), the intermediate domain deletion (Flag-MyD88(ΔID)), or the Toll-interleukin-1 receptor domain deletion (Flag-MyD88(ΔTIR)) (Fig. 4E).

Since MyD88 DD contains a hemi-immunoreceptor tyrosine-based activation motif (ITAM), the MyD88 hemi-ITAM deletion mutant (Flag-MyD88(ΔITAM)) and the MyD88 tyrosine 58 residue mutant (Flag-MyD88(Y58F)) were generated (Fig. S5C) and the interaction between Syk and these MyD88 mutants was further examined. Syk interacted with WT MyD88 but did not interact with either Flag-MyD88(ΔITAM) or Flag-MyD88(Y58F) mutants (Fig. 4F). Moreover, the phosphorylation of Syk and the NF-κB-mediated luciferase reporter gene activity induced by WT MyD88 were significantly suppressed by Flag-MyD88(ΔITAM) and Flag-MyD88(Y58F) mutants (Fig. 4G – H). These results indicate that phosphorylation of MyD88 at tyrosine 58 residue is required for interacting with Syk kinase.

### Src activated MyD88 by phosphorylation of the tyrosine 58 residue to interact with and activate Syk

The mechanism by which MyD88 is phosphorylated in macrophages was next investigated. To examine whether MyD88 is phosphorylated in macrophages, RAW264.7 cells were stimulated with LPS and MyD88 phosphorylation was evaluated. MyD88 was immediately phosphorylated by LPS at 1 min, and its phosphorylation gradually decreased after 1 min in the LPS-stimulated RAW264.7 cells (Fig. 5A). To examine whether MyD88 is phosphorylated at the tyrosine 58 residue in the hemi-ITAM, HEK293 cells were transfected with the constructs of WT Flag-MyD88, Flag-MyD88(ΔITAM), and Flag-MyD88(Y58F) mutants and MyD88 phosphorylation was further evaluated. WT MyD88 was phosphorylated, while neither Flag-MyD88(ΔITAM) nor Flag-MyD88(Y58F) mutants were phosphorylated (Fig. 5B).

**Figure 5.**
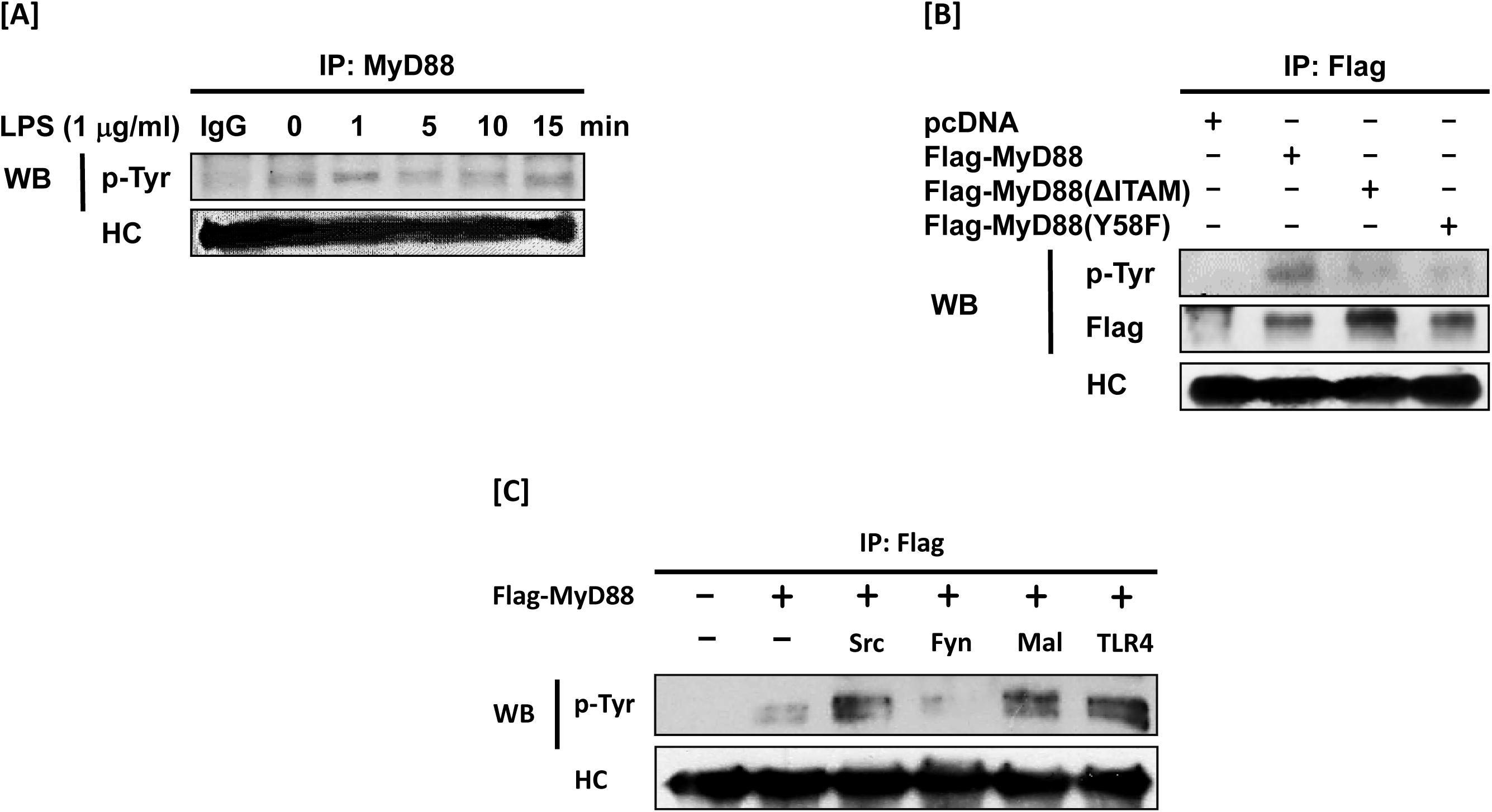
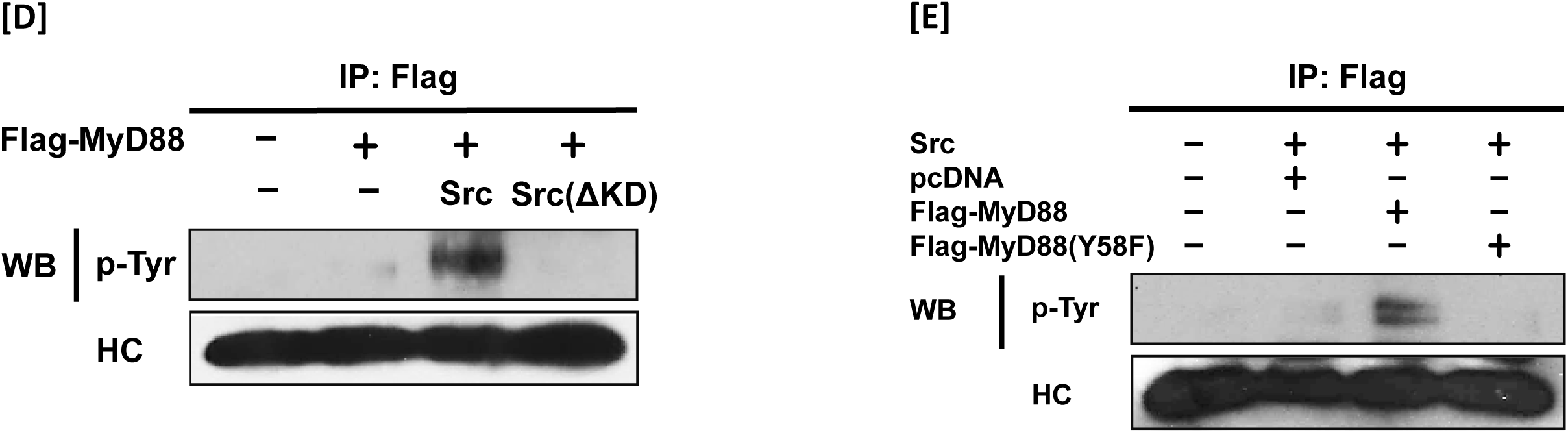
Src activated MyD88 by phosphorylation at the tyrosine 58 residue to interact with and activate Syk. (A) p-Tyr MyD88 in the whole cell lysates of RAW264.7 cells treated with LPS (1 μg/ml) for the indicated time was detected by immunoprecipitation and Western blot analysis. (B) p-Tyr MyD88 and Flag in the whole cell lysates of HEK293 cells transfected with either empty (pcDNA), Flag-MyD88, Flag-MyD88(ΔITAM) or Flag-MyD88(Y58F) plasmids for 48 h were detected by immunoprecipitation and Western blot analysis. (C) p-Tyr MyD88 in the whole cell lysates of RAW264.7 cells co-transfected with Flag-MyD88 and either empty (pcDNA), Src, Fyn, Mal, or TLR4 for 48 h was detected by immunoprecipitation and Western blot analysis. (D) p-Tyr MyD88 in the whole cell lysates of RAW264.7 cells co-transfected with Flag-MyD88 and either empty (pcDNA), Src, or Src(ΔKD) was detected by immunoprecipitation and Western blot analysis. (E) p-Tyr MyD88 in the whole cell lysates of RAW264.7 cells co-transfected with Src and either empty (pcDNA), Flag-MyD88, or Flag-MyD88(Y58F) plasmids for 48 h were detected by immunoprecipitation and Western blot analysis.

To identify the molecule that phosphorylates MyD88 in macrophages, RAW264.7 cells were transfected with TLR4, Mal (an adaptor interacting with the intracellular domain of TLR4), and the Src family kinases Src and Fyn. TLR4, Mal, and Src, but not Fyn, phosphorylated MyD88 (Fig. 5C). Src was phosphorylated by LPS faster than MyD88 from 15 to 30 sec, and its phosphorylation was notably inhibited beyond 30 sec in the LPS-stimulated RAW264.7 cells (Fig. S6). Therefore, Src-mediated MyD88 phosphorylation was further investigated. To confirm whether MyD88 is phosphorylated by Src, RAW264.7 cells were transfected with the WT Src and the Src kinase domain deletion mutant (Src(ΔKD)) and MyD88 phosphorylation was examined. MyD88 was phosphorylated by WT Src but not by Src(ΔKD) (Fig. 5D). Since MyD88 was phosphorylated at the tyrosine 58 residue by LPS (Fig. 5B), whether Src phosphorylates MyD88 at the tyrosine 58 residue was next examined, and it was found that Src did phosphorylate MyD88 at the tyrosine 58 residue (Fig. 5E).

### Src was activated by F-actin and Rac1

Finally, how Src is activated in macrophages was investigated. The formation of F-actin by actin polymerization was increased by LPS at 15 sec and dramatically decreased after 30 sec in RAW264.7 cells (Fig. 6A). To examine the role of actin polymerization on Src activation, RAW264.7 cells were treated with cytochalasin B (CytoB), an actin polymerization inhibitor (MacLean-Fletcher & Pollard, 1980), and Src phosphorylation was evaluated. CytoB clearly inhibited the actin polymerization at 15 and 30 sec in the LPS-stimulated RAW264.7 cells (Fig. S7). Src was highly phosphorylated by LPS from 15 to 30 sec, while the Src phosphorylation was markedly inhibited by CytoB in the LPS-stimulated RAW264.7 cells (Fig. 6B). Moreover, in accordance with the Src phosphorylation, Syk phosphorylation was also inhibited by CytoB in the LPS-stimulated RAW264.7 cells (Fig. 6C).

**Figure 6.**
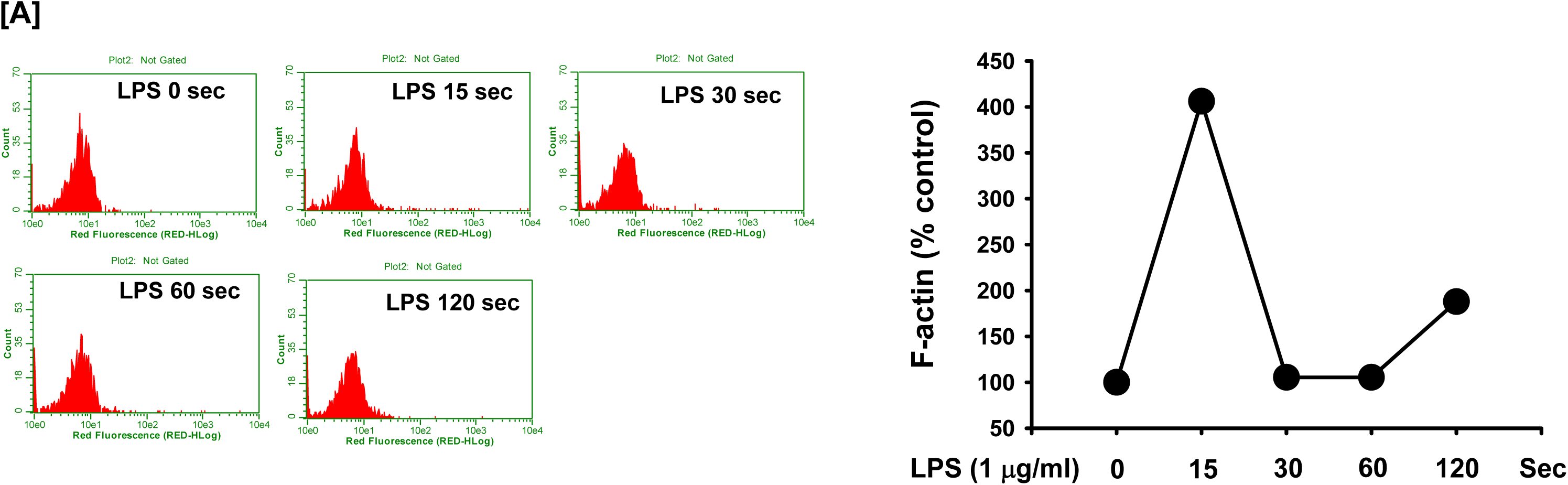
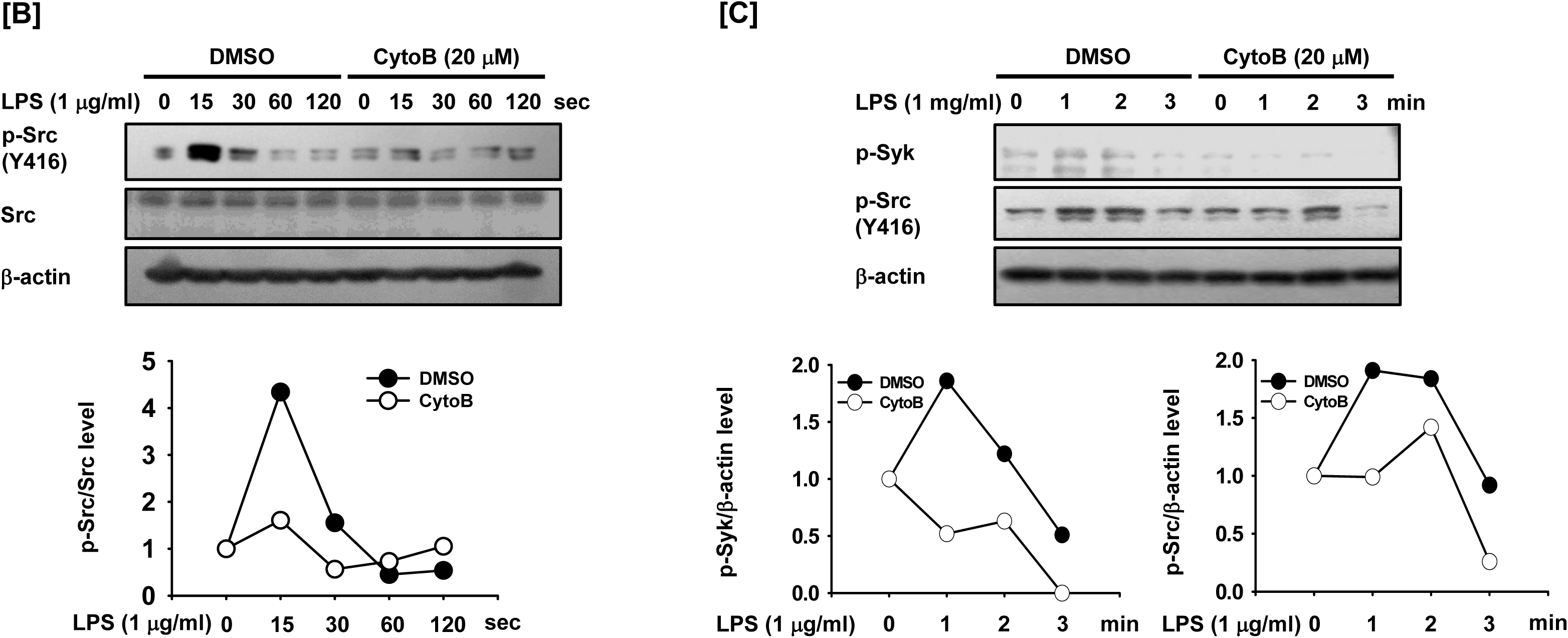
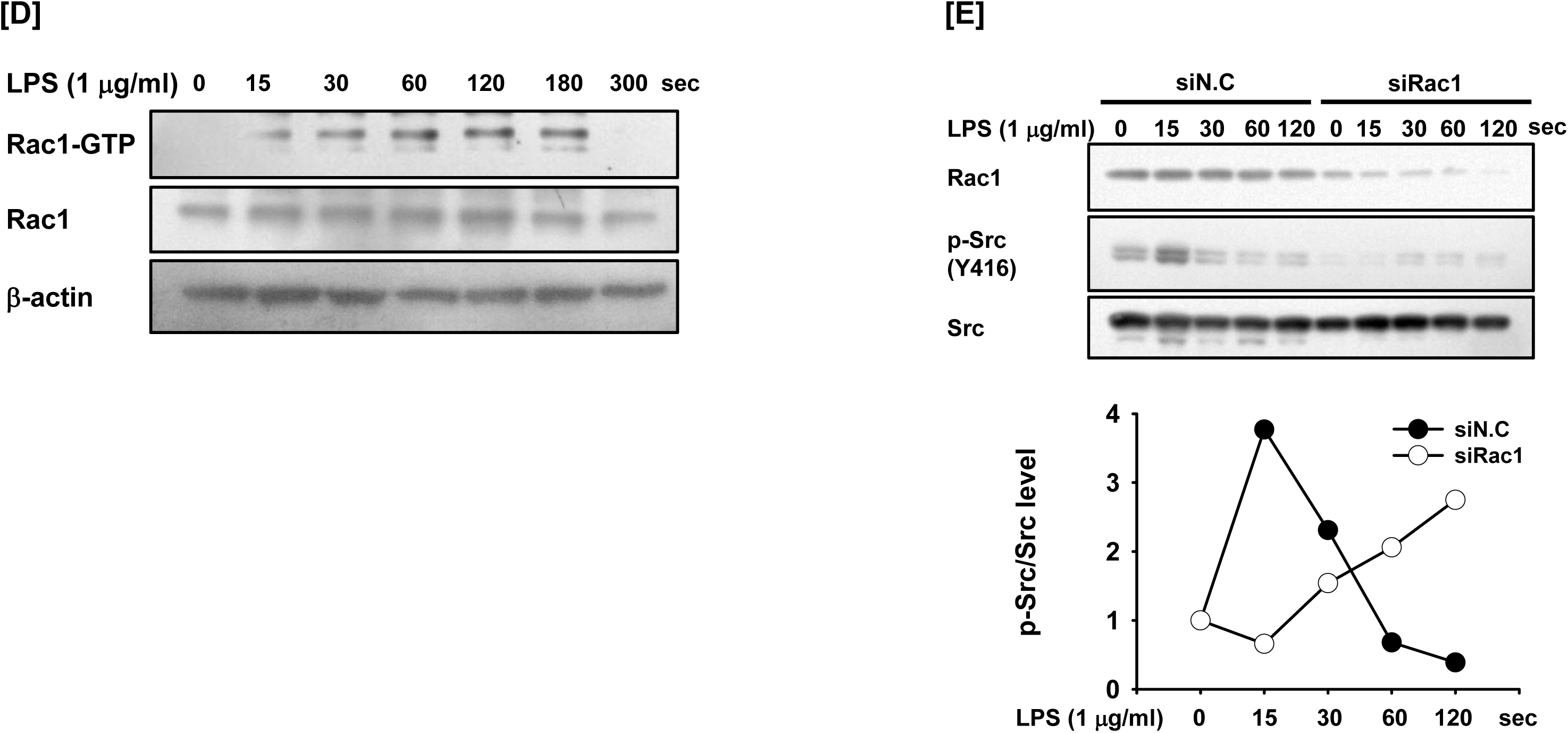
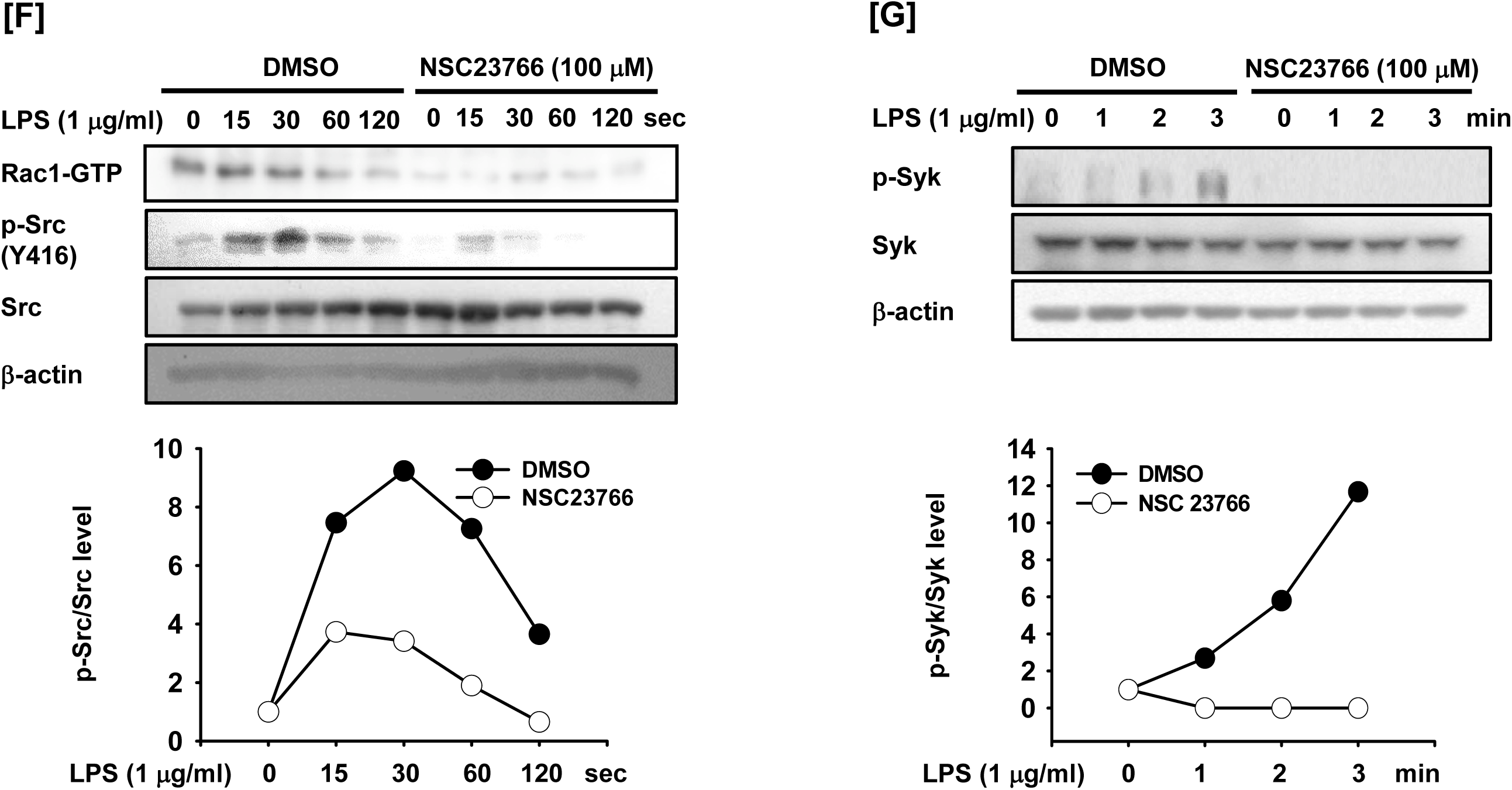
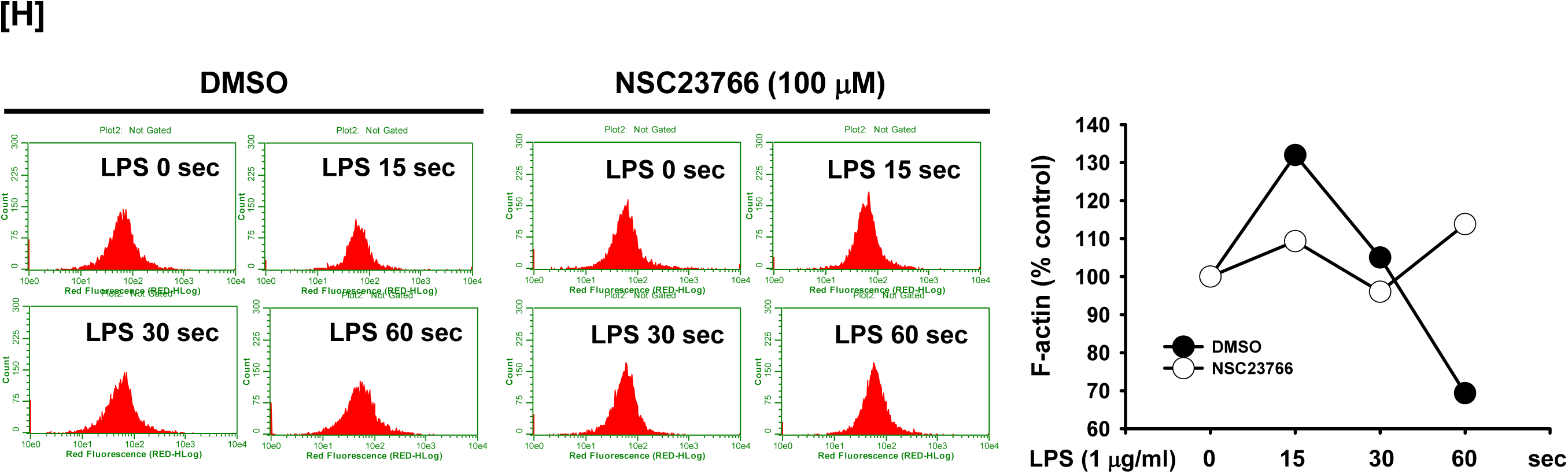
Src was activated by F-actin and Rac1. (A) RAW264.7 cells were treated with LPS (1 μg/ml) for the indicated time in the presence of Texas Red-X phalloidin (200 units/ml), and the fluorescence was determined by a flow cytometer and plotted. (B) Src and p-Src in the whole cell lysates of RAW264.7 cells incubated with vehicle (DMSO) or CytoB (20 μM), followed by the treatment of LPS (1 μg/ml) for the indicated time were detected by Western blot analysis, and the p-Src/Src level was plotted. (C) p-Syk and p-Src in the whole cell lysates of RAW264.7 cells incubated with vehicle (DMSO) or CytoB (20 μM), followed by the treatment of LPS (1 μg/ml) for the indicated time were detected by Western blot analysis, and the p-Syk/β-actin and p-Src/β-actin levels were plotted. (D) Rac1-GTP and Rac1 in the whole cell lysates of RAW264.7 cells treated with LPS (1 μg/ml) for the indicated time were detected by Western blot analysis. (E) Rac1, Src, and p-Src in the whole cell lysates of RAW264.7 cells transfected with siN.C or siRac1 for 24 h, followed by the treatment of LPS (1 μg/ml) for the indicated time were detected by Western blot analysis, and the p-Src/Src level was plotted. (F) Rac1-GTP, Src, and p-Src in the whole cell lysates of RAW264.7 cells incubated with vehicle (DMSO) or NSC23766 (100 μM), followed by the treatment of LPS (1 μg/ml) for the indicated time were detected by Western blot analysis, and the p-Src/Src level was plotted. (G) Rac1-GTP, Syk, and p-Syk in the whole cell lysates of RAW264.7 cells incubated with vehicle (DMSO) or NSC23766 (100 μM), followed by the treatment of LPS (1 μg/ml) for the indicated time were detected by Western blot analysis, and the p-Syk/Syk level was plotted. (H) RAW264.7 cells incubated with vehicle (DMSO) or NSC23766 (100 μM) were treated with LPS (1 μg/ml) for the indicated time in the presence of Texas Red-X phalloidin (200 units/ml), and the fluorescence was determined by a flow cytometer and plotted.

The role of the Rho GTPase family member Rac1 on Src activation was next examined. Rac1-GTP levels were increased by LPS in RAW264.7 cells (Fig. 6D), and inhibition of Rac1 expression by Rac1 siRNA (siRac1) notably suppressed Src phosphorylation in the LPS-stimulated RAW264.7 cells (Fig. 6E). Inhibition of Rac1 activation by its selective inhibitor NSC23766 also suppressed the phosphorylation of Src (Fig. 6F) and Syk (Fig. 6G) in the LPS-stimulated RAW264.7 cells. Moreover, inhibition of Rac1 activation by NSC23766 suppressed the actin polymerization (Fig. 6H), suggesting that Rac1 acts as an upstream molecule to regulate actin polymerization in Src regulatory mechanism.

## Discussion

Previous studies have demonstrated that Syk and MyD88 are involved in the inflammatory response through activation of the inflammatory signaling pathways in macrophages. However, despite a number of studies, their roles and the corresponding molecular mechanism are still largely unidentified. Moreover, the functional relationships between key molecules for inflammatory responses are poorly understood. Therefore, the present study explored the roles and the functional cooperation of Syk and MyD88 as well as the underlying molecular mechanisms during inflammatory responses in macrophages.

Although ROS/RNS generation and phagocytosis are critical processes of the inflammatory response in macrophages, understanding of the molecular mechanism by which ROS/RNS and phagocytosis are generated in the macrophage-mediated inflammatory response is still very limited. However, it has been demonstrated that stimulation of macrophages by PAMPs such as LPS induces ROS/RNS generation and phagocytosis (Lucas & Maes, 2013), and that many intracellular signaling molecules, including Syk, are activated during the LPS-stimulated inflammatory response (Byeon et al, 2012; Liu et al, 2010; Yang et al, 2014; Yi et al, 2014).

In this study, a Syk-deficient condition was generated in RAW264.7 cells in three ways: CRISPR/Cas9, siSyk, and a selective Syk inhibitor, Pic. Regardless of the Syk inhibition method, LPS-induced ROS/RNS generation and phagocytic activity were significantly decreased in the Syk-deficient RAW264.7 cells from 10 to 30 min (Fig. 1), indicating that Syk is a critical player to induce the ROS/RNS generation and phagocytic activity of macrophages rapidly in 30 min during the inflammatory response.

The generation of ROS/RNS performs a variety of biological functions, including signal transduction and differentiation in macrophages, as well as pathogen killing through respiratory bursts (Covarrubias et al, 2013). In this study, it was confirmed that ROS/RNS generated from 10 minutes after E.coli treatment were closely involved in phagocytosis (Fig. 1K and L). Interestingly, ROS/RNS generated by LPS treatment increased the N-Tyr of SOC1, resulting in phagocytosis (Fig.2I). This result implies that the generated ROS/RNS act as a signaling molecule to induce phagocytosis by regulating the activity of SOCS1. SOCS1 overexpression decreased phagocytic activity, and SOCS1 inhibition by its specific siRNA (siSOCS1) increased phagocytic activity in the LPS-stimulated RAW264.7 cells, strongly suggesting that SOCS1 is a negative regulator of phagocytic activity in inflammatory responses. Tyrosine nitrification studies of manganese- dependent superoxide dismutase (MnSOD) and cytochrome c show that such modifications can eventually lead to loss-of-function (MnSOD) or gain-of-function (cytochrome c) through structural modification of the protein (Abriata et al, 2009; Yamakura et al, 1998). Eventually, since LPS increases the N-Tyr of SOCS1, a negative regulator (Fig.S3), it can be assumed that N-Tyr of SOCS1 induced by ROS/RNS generation will inactivate SOCS1.

To understand how Syk induces ROS/RNS generation and its molecular mechanism, the role of Syk on the expression of ROS/RNS-generating enzymes in macrophages was examined. Several ROS/RNS-generating enzymes have been identified. iNOS catalyzes L-arginine to produce NO (Suschek et al, 2004), and NOX1 produces ROS/RNS by oxidizing NADPH (Makhezer et al, 2019). AOC3 catalyzes oxidative deamination of primary amines, producing H_2_O_2_ (Lyles, 1996). AOX1 produces ROS/RNS by oxidizing various aldehydes (Kundu et al, 2007), and XDH also generates ROS/RNS by oxidizing ethanol (Castro et al, 2001). mRNA expression of these enzymes was increased by LPS in RAW264.7 cells, but interestingly, mRNA expression of iNOS, NOX1, AOX1, and XDH was rapidly increased at 10 min, whereas AOC3 mRNA expression was increased at 30 min in the LPS-stimulated RAW264.7 cells (Fig. 2A), implying that although these are all ROS/RNS-generating enzymes, their modes of action to induce inflammatory responses in macrophages might be different. mRNA expression of these enzymes induced by LPS was markedly decreased in the Syk^-/-^ and Syk- deficient RAW264.7 cells compared to control WT cells (Fig. 2B – D), indicating that Syk induces ROS/RNS generation by upregulating the expression of ROS/RNS-generating enzymes in macrophages during inflammatory responses.

Moreover, inhibition of Syk markedly suppressed the NF-κB signaling pathway (Fig. 3A – F). In particular, Syk appears to play an important role in phosphorylation of IκBα at early time point (5 min) under LPS treatment. Interestingly, only MyD88 was involved in IκBα phosphorylation at 5 minutes, among MyD88 and TRIF which are adaptors that interact with the intracellular domain of TLR4 (Fig. 3G and S4E). It has been also reported that MyD88 and Syk are key players to activate the NF-κB inflammatory signaling pathway in macrophages (Yi et al, 2014), therefore we expected MyD88 to be an upstream molecule of Syk kinase under LPS stimulation condition. In accordance with the results of the Syk-deficient RAW264.7 cells, mRNA expression of ROS/RNS-generating enzymes, ROS/RNS generation, and phagocytic activity induced by LPS were notably decreased in the MyD88-/- RAW264.7 cells compared to control WT cells (Fig. S4F - I). Suppression of Syk dramatically inhibited the NF-κB luciferase reporter gene activity induced by MyD88, strongly indicating that Syk and MyD88 are key players that induce ROS/RNS generation and ROS/RNS-mediated phagocytic activity by activating the NF-κB signaling pathway in macrophages during the inflammatory response.

As discussed earlier, Syk and its upstream molecule, MyD88, induced ROS/RNS generation and the phagocytic activity of macrophages during LPS-induced inflammatory responses (Figs. 1 and 2), however, how they cooperate in these inflammatory processes was still unclear. Therefore, the functional relationship between Syk and MyD88 was next examined in the LPS-stimulated RAW264.7 cells. MyD88 activated Syk (Fig. 4A), and MyD88-induced activation of NF-κB was abolished by a defect of the Syk kinase domain (Fig. 4B), indicating that MyD88 is an upstream regulator of Syk-mediated NF-κB activation. The question of how MyD88 cooperates with Syk for Syk activation was raised. ITAM, which has the consensus sequence of YxxI/Lx_6-12_YxxI/L, is a conserved phosphorylation module found in a large number of adaptors and receptors and engages in a variety of biological processes and functions (Bezbradica & Medzhitov, 2012). Receptor ligation induces the tyrosine phosphorylation of ITAM by the intracellular kinase families, which in turn leads to the recruitment, interaction, and activation of Syk (Abram & Lowell, 2007). Interestingly, hemi-ITAM, a single-copy of ITAM, was found in MyD88 (YLEI; 58 – 61); therefore, the molecular interaction between Syk and MyD88 was examined. Syk interacted with MyD88 in RAW264.7 cells and the peritoneal macrophages isolated from mice (Fig. 4C). To examine which domains of Syk and MyD88 are crucial for their molecular interaction, domain deletion mutants of Syk (Fig. S5A) and MyD88 (Fig. S5B) and their interaction were examined. MyD88 interacted with Syk(ΔKD) but did not interact with Syk(ΔSH2-N) and Syk(ΔSH2-C) (Fig. 4D), indicating that the two SH2 domains of Syk are critical but the kinase domain of Syk is dispensable for interaction with MyD88. In addition, Syk interacted with only WT MyD88, and none of the MyD88 mutants interacted with Syk (Fig. 4E). This result could be explained by two possibilities. The first is that all three domains of MyD88 are critical for the interaction with Syk, and the second is that deleting any one of the three domains could induce a significant conformational change, resulting in failure to interact with Syk. To clarify these possibilities and to examine other possibilities, more studies need to be conducted. Since ITAM is the site for Syk interaction and the tyrosine is one of the phosphorylated amino acids for protein functions, the interaction of Syk with MyD88(ΔITAM) and the substitution mutant of tyrosine in ITAM to phenylalanine, MyD88(Y58F), was further examined. Syk did not interact with either MyD88(ΔITAM) or MyD88(Y58F) (Fig. 4F), indicating that hemi-ITAM in MyD88 is essential for the interaction with Syk and the tyrosine residue in the ITAM of MyD88 is a critical amino acid for the interaction with Syk. In addition, the activation of Syk and the NF-κB-luciferase activity induced by WT MyD88 was markedly suppressed by MyD88(ΔITAM) and MyD88(Y58F) (Fig. 4G – H). These results suggest that the interaction of Syk with its upstream molecule MyD88 is crucial for the activation of Syk and the consequent NF-κB signaling pathway in macrophages during the inflammatory responses and that ITAM and the tyrosine 58 residue in the ITAM of MyD88 play an essential role for Syk-MyD88 interaction.

Receptor-ligand interaction induces the phosphorylation of ITAM in the receptors and adaptors to initiate the signal transduction cascades in cells (Abram & Lowell, 2007), which raised the question whether MyD88 is phosphorylated at the tyrosine 58 residue in the hemi-ITAM for its activation during the macrophage-mediated inflammatory responses. MyD88 was immediately phosphorylated by LPS around 1 min (Fig. 5A), and the tyrosine 58 residue was identified as the main amino acid phosphorylated (Fig. 5B). Several intracellular molecules, such as Src family kinases and TLR4 adaptors, have been identified as very upstream molecules immediately activated by PAMPs to induce the inflammatory responses in macrophages (Byeon et al, 2012; Yang et al, 2014; Yi et al, 2014; Yu et al, 2012), and hence, which molecule is responsible for the phosphorylation of MyD88 at the tyrosine 58 residue was examined. To identify the upstream molecules that phosphorylate MyD88, MyD88 was co-transfected with TLR4, Mal, and the Src family kinases Src and Fyn. TLR4 and Mal induced MyD88 phosphorylation (Fig. 5C), which is an inevitable result since MyD88 activation is induced by TLR4 and its adaptors (Yi et al, 2014). Interestingly, among the Src family kinases, Src, but not Fyn, induced MyD88 phosphorylation (Fig. 5C), indicating that despite belonging to the same family, they have distinct roles and not all Src family kinases stimulate MyD88 phosphorylation and MyD88-induced signal transduction cascades in macrophage-mediated inflammatory responses. Therefore, Src-induced MyD88 phosphorylation was further examined. Src was phosphorylated by LPS quickly at 15 to 30 sec, lasting up to 120 sec (Fig. S6), indicating that Src is activated by phosphorylation faster than MyD88 phosphorylated by LPS at around 1 min (Fig. 5A) and that Src might be an upstream regulator of MyD88. In accordance with the previous result (Fig. 5C), Src induced MyD88 phosphorylation, while MyD88 phosphorylation induced by Src was abolished by Src(ΔKD) (Fig. 5D). Moreover, Src-induced MyD88 phosphorylation was also abolished by MyD88(Y58F). These results suggest that Src is an upstream activator of MyD88 by phosphorylating it at the tyrosine 58 residue.

The mechanism by which Src is activated is well established. Src is activated by dephosphorylation at the 527 tyrosine residue by various tyrosine phosphatases and the subsequent autophosphorylation at the 416 tyrosine residue in the kinase domain. However, knowledge of which molecules are involved in and responsible for Src phosphorylation is still lacking. Therefore, how Src is activated in macrophages during the inflammatory response was finally examined. A previous study demonstrated that polymerization of globular actin to F-actin is a key process in the inflammatory response (Du et al, 2012). Another study reported that disruption of the F-actin cytoskeleton structure induced Src deactivation by suppressing the formation of a signaling complex consisting of Src and p85/PI3K (Kim et al, 2013), suggesting that the inflammatory signals induce the formation of F-actin, and the F-actin subsequently activates Src, leading to the induction of the inflammatory signal transduction cascades in macrophages. To examine this, RAW264.7 cells were stimulated by LPS, and F-actin formation was evaluated. LPS immediately induced the formation of F-actin at 15 sec, and the formation of F-actin was decreased after 15 sec in RAW264.7 cells (Fig. 6A). CytoB, a selective inhibitor of actin filament formation, reduced the formation of F-actin (Fig. S7) and markedly suppressed the Src phosphorylation induced by LPS in RAW264.7 cells (Fig. 6B), leading to the subsequent inhibition of the Syk phosphorylation in the LPS-stimulated RAW264.7 cells (Fig. 6C). These results suggest that actin polymerization is immediately induced in macrophages during the inflammatory responses, resulting in the activation of Src to transduce downstream inflammatory signals. Although F-actin activates Src by phosphorylation, F-actin is not a kinase; therefore, F-actin might indirectly activate Src phosphorylation, e.g. by inhibiting the formation of an inflammatory signaling complex composed of Src and p85/PI3K, as previously discussed (Kim et al, 2013), or by activating other Src-phosphorylating kinases. Further studies in this regard are highly needed. Rac1, a small G-protein belonging to the Rho protein family, is implicated in various biological processes, such as fibrosis, apoptosis, gene regulation, and tumorigenesis (Bid et al, 2013; Nagase & Fujita, 2013). Rac1 is also reported to play roles as an upstream regulator in inducing inflammatory responses by increasing ROS/RNS generation and activating the NF-κB signaling pathway in response to a number of stimulations (Cuadrado et al, 2014; Nagase & Fujita, 2013). Therefore, the role of Rac1 in Src activation was further examined. Similar to F-actin, Rac1 was immediately activated by LPS from 15 sec to 180 sec (Fig. 6D), and inhibition of Rac1 suppressed phosphorylation of Src and Syk induced by LPS in RAW264.7 cells (Fig. 6E - G). These results indicate that Rac1, like F-actin, is an upstream activator of Src in macrophages during the inflammatory response. Similar to the previous observation that Rho GTPases, including Rac1, induce actin polymerization (Hall, 1998; Tata et al, 2014), inhibition of Rac1 activation by NSC23766 suppressed the formation of F-actin (Fig. 6H). This result suggests that Rac1 induces polymerization of actin, leading to the activation of Src in macrophages during inflammatory responses.

In conclusion, the current study demonstrated the roles of Syk and the underlying molecular mechanisms in macrophage-mediated inflammatory responses. Upon LPS stimulation, Syk induced ROS/RNS generation by upregulating the expression of ROS/RNS-generating enzymes as well as the phagocytic activity of macrophages by suppressing the negative regulator of phagocytosis, SOCS1, via nitration of SOCS1 in 10 to 30 min. Syk was activated by the molecular interaction with its upstream adaptor, MyD88, and the 58 tyrosine residue in the hemi-ITAM of MyD88 was identified as a critical site for their interaction. Inflammatory stimulation by LPS immediately induced F-actin formation and Rho-GTPase Rac1 activation from 15 sec, which activated Src at 15 to 30 sec, subsequently activating MyD88 by phosphorylation at the 58 tyrosine residue in 1 min in macrophages during the inflammatory response, as described in Figure 7. Taken together, our observations contribute to the understanding of the meaning and significance of the early activation of the MyD88-Syk axis and provide the mechanistic basis of these events in macrophage-mediated inflammatory responses. Moreover, since the MyD88-Syk axis is rapidly activated in macrophages during inflammatory responses, selective targeting of both or one of these molecules is a potentially promising strategy to prevent and treat inflammatory and autoimmune diseases.

**Figure 7.**
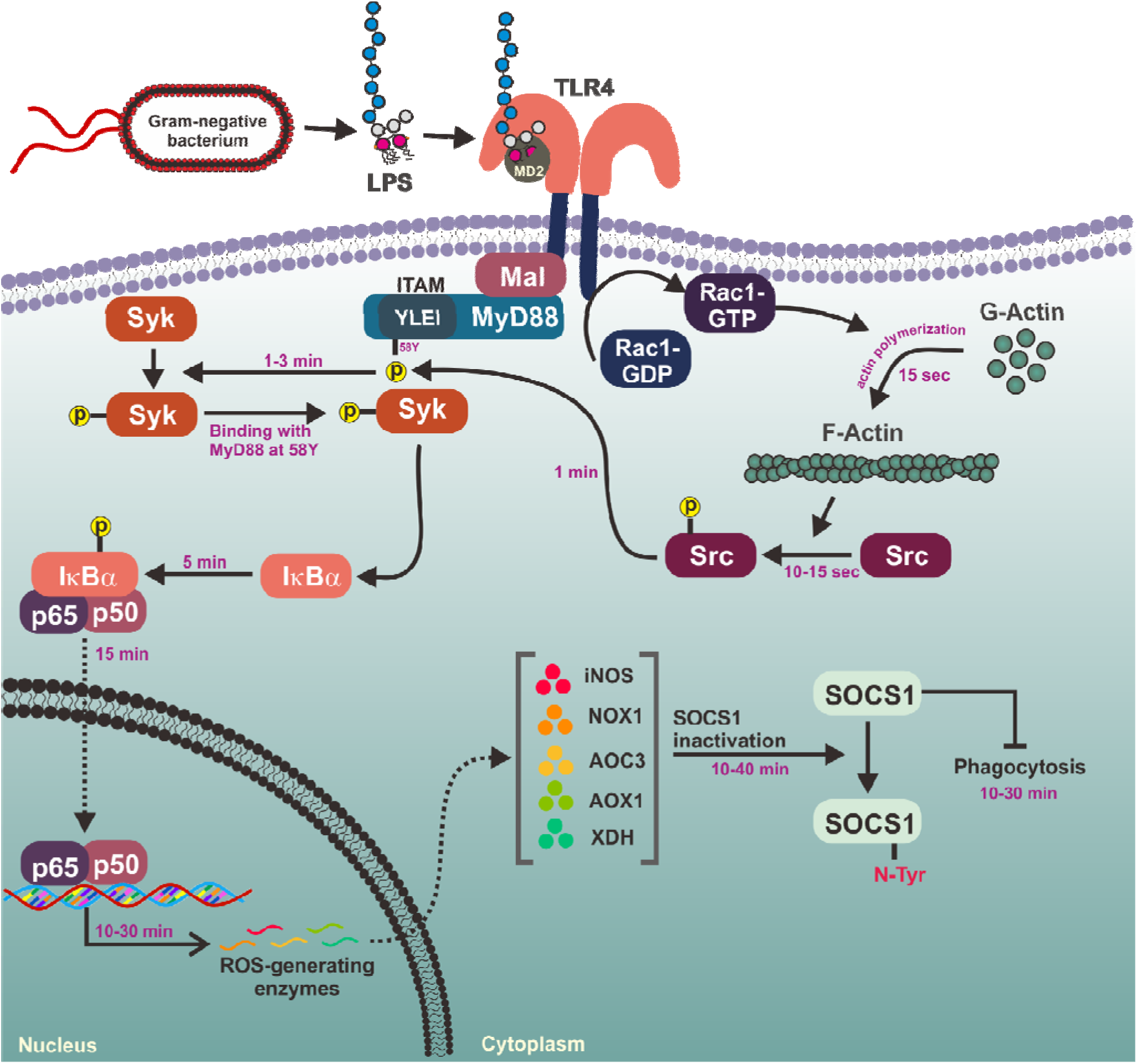
Schematic summary demonstrating the role of MyD88-Syk axis in the macrophage-mediated inflammatory responses

## Materials and Methods

### Materials

Normal C57BL/6 mice (male, 6 weeks, 20-25 g) were purchased from Daehan Bio Link Co., Ltd. (Osong, Korea). Knockout mice without MyD88 or TRIF were obtained from Prof. Narayanan Parameswaran (Department of Physiology, Michigan State University, Michigan, USA). Mouse pellet diet was purchased from Samyang (Daejeon, Korea). Roswell Park Memorial Institute 1640 (RPMI 1640) cell culture medium, Dulbecco’s Modified Eagle’s medium (DMEM), fetal bovine serum (FBS), streptomycin, penicillin, L-glutamine, phosphate-buffered saline (PBS), Lipofectamine^®^ 2000, mounting media, TRIZOL^TM^ Reagent, MuLV reverse transcriptase, *pfu* DNA polymerase, DH5α competent cells, polyvinylidene difluoride (PVDF) membrane, enhanced chemiluminescence (ECL) reagent, and Texas Red-X phalloidin were purchased from Thermo Fisher Scientific (Waltham, MA, USA). LentiCRISPRv2 vector was purchased from Addgene (Watertown, MA, USA). Puromycin dihydrochloride, lipopolysaccharide (LPS, *Escherichia coli* 0111:B4), dihydrorhodamine 123 (DHR 123), FITC-E.coli, piceatannol (Pic), bovine serum albumin (BSA), α-tocopherol, resveratrol, tumor necrosis factor-α (TNF-α), phorbol-12-myristate-13-acetate (PMA), luciferin, N(G)-nitro-L-arginine methyl ester (L-NAME), sodium nitroprusside (SNP), sodium dodecyl sulfate (SDS), and CytoB were purchased from Sigma Chemical Co. (St. Louis, MO, USA). LentiCRISPRv2 plasmid, NF-κB-luciferase reporter gene plasmid, and β-galactosidase plasmid were purchased from Addgene (Cambridge, MA, USA). Antibodies specific for LAMP-1 (ab25245), FITC (sc-69872), phosphor (p)-IκBα (CST 9246), IκBα (CST 9247), p65 (CST 8242), p50 (CST 12540), N-Tyr (ab24497), p-tyrosine (Upstate 05-321), p-serine (SC 81514), p-threonine (CST 9381), SOCS1 (ab9870), p-STAT1 (CST 9167), STAT1 (CST 9172), p-Syk (CST 2711), Syk (CST 2712), MyD88 (CST 4283), p-Src(Y416) (CST 2101), Src (CST 2109), Fyn (CST 4023), Mal (sc-390687), TLR4 (CST 14358), Rac1-GTP (NB 26903), Rac1 (CST 4651), p-p85 (CST 4228), p85 (CST 4292), TRIF (CST 13077), Flag (CST 8146), Myc (CST 2276), β-actin (CST 4967), and lamin A/C (CST 4777) used for Western blot analysis, immunoprecipitation, and immunofluorescence staining were purchased from Abcam (Cambridge, UK), Cell Signaling Technology (Beverly, MA, USA), Santa Cruz Biotechnology, Inc. (Dallas, TX, USA), Upstate Biotechnology (Waltham, MA, USA), and NewEast Biosciences (Malvern, PA, USA). QIAprep Spin Miniprep Kit was purchased from Qiagen, Inc. (Germantown, MD, USA). Automated DNA sequencing of mutant plasmids was performed at Bionics (Seoul, Korea). siRNAs for Syk and SOCS1 were purchased from Genolution (Seoul, Korea). sgRNAs for Syk, MyD88, and TRIF and primers used for quantitative real-time PCR were synthesized at Macrogen Inc. (Seoul, Korea). SYBR Premix ex Taq was purchased at Takara Bio Inc. (Shiga, Japan). Protein A-coupled Sepharose^®^ beads were purchased at GE Healthcare Life Sciences (Marlborough, MA, USA).

### Mice, husbandry, and preparation of peritoneal macrophages

C57BL/6 mice (male, 6 weeks, 20 – 25 g) housed in a plastic cage at 22°C in a 12 h light-dark cycle were fed pelleted mouse diet and tap water *ad libitum*. Animal care and the study using the mice were conducted according to the guidelines of and in compliance with protocols approved by the Institutional Animal Care and Use Committee at Sungkyunkwan University. Peritoneal exudates were obtained from C57BL/6 male mice by lavage 4 days after intraperitoneal injection of 1 ml sterile 4% thioglycolate broth (Difco Laboratories, Detroit, MI, USA), as previously reported (Kim et al, 2019). After the exudates were washed with RPMI1640 medium containing 2% FBS, peritoneal macrophages (1×10^6^ cells/ml) were plated in 100mm tissue culture dishes for 4 h at 37 °C in a 5% CO2 humidified atmosphere.

### Cells culture and transfection

RAW264.7 and human embryonic kidney 293 (HEK293) cells were purchased from American Type Culture Collection (Rockville, MD, USA). Peritoneal macrophages from C57BL/6 mice were prepared as described previously (Kim et al, 2018). RAW264.7 cells and peritoneal macrophages were culture in RPMI 1640 medium, and HEK293 cells were cultured in DMEM. RPMI 1640 media and DMEM were supplemented with 10% FBS, streptomycin (100 mg/ml), penicillin (100 U/ml), and L-glutamine (2 mM) at 37°C in a 5% CO_2_ humidified incubator. Mycoplasma contamination was regularly tested using the BioMycoX Mycoplasma PCR Detection Kit (CellSafe, Seoul, Korea). RAW264.7 and HEK293 cells were transfected with either expression constructs, siRNA (Supplementary Table 1), or sgRNA (Supplementary Table 2) constructs for 48 h using Lipofectamine^®^ 2000 according to the manufacturer’s instructions.

### Generation of Syk^-/-^, MyD88^-/-^, and TRIF^-/-^ RAW264.7 cells by CRISPR/Cas9 system

sgRNA sense and anti-sense oligos of Syk, MyD88, and TRIF were synthesized, and their sgRNA constructs were generated by inserting each sgRNA hybrid into a lentiCRISPRv2 vector, as described previously (Shalem et al, 2014; Yi et al, 2018). RAW264.7 cells were transfected with either Syk, MyD88, or TRIF sgRNA construct using Lipofectamine^®^ 2000 according to the manufacturer’s instructions, and the transfected RAW264.7 cells were selected using puromycin (1.0 mg/ml) in RPMI 1640 medium containing 10% FBS until all non-transfected RAW264.7 cells were dead.

### ROS/RNS generation assay

RAW264.7 cells treated with LPS (1 μg/ml) at 37°C for the indicated time were treated with DHR 123 (20 μM) at 37°C for 20 min. After washing the cells three times with PBS, the cells were resuspended in FACS buffer (2% BSA, 0.1% sodium azide in PBS) and the fluorescence of the cells was determined using a flow cytometer.

### Phagocytosis assay

RAW264.7 cells were treated with FITC-E.coli (10 μg/ml) for the indicated time. After washing the cells three times with PBS, the cells were resuspended in FACS buffer and the fluorescence of the cells was determined using a fluorescence microplate reader and a flow cytometer.

### Confocal microscopy

RAW264.7 cells were transfected with either control scramble siRNA (siN.C) or Syk siRNA (siSyk) for 24 h. RAW264.7 cells were treated with either vehicle (DMSO), α-tocopherol (100 mM), or resveratrol (50 mM) for 12 h. The RAW264.7 cells were treated with FITC-E.coli (10 μg/ml) for the indicated time and fixed with formaldehyde (4%) for 10 min at room temperature, washed with PBS three times for 5 min each, and permeabilized with Triton X-100 (1%) for 5 min. After washing the cells, BSA was added to the cells for blocking, and the cells were incubated with the primary antibodies specific for LAMP-1 or FITC, followed by secondary antibody incubation. After washing the cells with PBS three times for 5 min each, the cells were covered with mounting media (20 μl) and a cover glass. The cells were then analyzed by confocal microscopy.

### Quantitative real-time polymerase chain reaction (PCR)

Total RNA was extracted using TRIZOL^TM^ Reagent according to the manufacturer’s instructions and immediately stored at −70°C until use. 1 μg of the total RNA was used for synthesizing cDNA using MuLV reverse transcriptase, and the cDNA was used for quantitative real time PCR using SYBR Premix ex Taq according to the manufacturer’s instructions. Primer sequences used for quantitative real-time PCR are listed in Supplementary Table 3.

### Luciferase reporter gene assay

HEK293 cells transfected with NF-κB-Luciferase reporter gene constructs and β-galactosidase-expressing constructs were treated or co-transfected with the indicated reagents or constructs using Lipofectamine^®^ 2000 according to the manufacturer’s instructions. The cells were lysed by repeating freezing at −70°C and thawing at room temperature three times. Luciferase reporter gene activity was determined by adding luciferase substrate, luciferin, to the cell lysates using a luminometer. All luciferase reporter gene activities were normalized to the β-galactosidase activity.

### Preparation of whole cell and nuclear lysates

For preparation of whole cell lysates, RAW264.7 and HEK293 cells were lysed with ice-cold buffer A (20 mM Tris-HCl, pH 7.4, 2 mM EDTA, 2 mM EGTA, 50 mM glycerol phosphate, 1 mM DTT, 2 µg/ml aprotinin, 2 µg/ml leupeptin, 1 μg/ml pepstatin, 50 μM PMSF, 1 mM benzamide, 2% Triton X-100, 10% glycerol, 0.1 mM sodium vanadate, 1.6 mM pervanadate, and 20 mM NaF) on ice for 30 min, followed by centrifugation at 12,000 rpm at 4°C for 5 min. The supernatant was collected as whole cell lysates and stored at −20°C until use.

For preparing nuclear lysates, RAW264.7 cells were lysed with ice-cold lysis buffer (10 mM HEPES pH 7.8, 1.5 mM MgCl_2_, 10 mM KCl, 0.5 mM DTT, 0.5% NP-40, 0.1 mM PMSF, 2 μg/ml leupeptin, and 2 μg/ml aprotinin) on ice for 10 min, followed by centrifugation at 3,000 rpm at 4°C for 10 min, and the supernatant was discarded. The pellet was resuspended in ice-cold buffer B (5 mM HEPES, 1.5 mM MgCl2, 0.2 mM EDTA, 0.5 mM DTT, 26% glycerol (v/v), 300 mM NaCl, pH 7.9), followed by sonication for 10 sec on ice, and the lysates were centrifuged at 20,000 g at 4°C for 30 min. The supernatant was collected as nuclear lysates and stored at −20°C until use.

### Western blot analysis and immunoprecipitation

For Western blot analysis, whole cell and nuclear lysates were subjected to SDS polyacrylamide gel electrophoresis and transferred on PVDF membranes. The membranes were blocked by blocking buffer (5% BSA in Tris-buffered saline) at room temperature for 1 h and incubated with primary antibodies specific for each target at room temperature for 1 h. After washing the membranes with washing buffer (Tris-base, NaCl, 0.1 % Tween 20, pH 7.6) three times for 10 min each, the membranes were incubated with HRP-linked secondary antibodies at room temperature for 1 h. After washing three times, the target proteins were visualized by the ECL reagent.

For immunoprecipitation, whole cell lysates (500 μg exogenous and 1 mg endogenous protein) were pre-cleared using 10 μl of protein A-coupled Sepharose^®^ beads (50% v/v) at 4°C for 1 h, and the pre-cleared whole cell lysates were incubated with primary antibodies specific for each target at 4°C overnight with gentle rotation. The immunocomplexes were then incubated with protein A-coupled Sepharose^®^ beads (50% v/v) at 4°C for 4 h with gentle rotation, and the supernatant was removed from the beads by centrifugation. The beads were washed with ice-cold buffer A five times and boiled for 5 min in protein sample buffer (50 mM Tris-HCl pH 6.8, 2% SDS, 10% glycerol, 1% β-mercaptoethanol, 12.5 mM EDTA, 0.02% bromophenol blue). Immunoprecipitates were analyzed by Western blot to detect target proteins.

### Construction of Syk and MyD88 mutants

Domain deletion mutants of Syk (ΔSH2-N, ΔSH2-C, and ΔKD) and MyD88 (ΔDD, ΔID, ΔTIR, and ΔITAM) and the tyrosine 58 residue point mutant (Y58F) were generated by site-directed mutagenesis. Mutant plasmids were generated by PCR using *pfu* DNA polymerase with mutant primers, and PCR conditions were as follow: pre-denaturation (95°C, 30 sec) followed by 18 cycles of denaturation (95°C, 30 sec), annealing (55°C, 1 min), and extension (68°C, 1 min/kb). The PCR-amplified mutant plasmids were transformed into DH5α competent cells, and the transformed DH5α competent cells were grown on LB agar plates containing ampicillin (100 μg/ml) at 37°C overnight. Mutant plasmids were prepared from the DH5α competent cells using the QIAprep Spin Miniprep Kit according to the manufacturer’s instructions, and mutagenesis was confirmed by automated DNA sequencing.

### Actin polymerization assay

RAW264.7 cells were treated with LPS and Texas Red-X phalloidin (200 units/ml) in the absence or presence of CytoB (20 μM) for the indicated time. After washing the cells three times with PBS, the cells were resuspended in FACS buffer, and fluorescence of the cells was determined using a flow cytometer.

### Statistical analysis

The data presented in this study are expressed as the mean ± standard deviation (SD) of at least three independent experiments. For statistical comparison, all results were analyzed by either analysis of variance (ANOVA) or the Mann-Whitney test, and a *P* value less than 0.05 was considered statistically significant (\**P* < 0.05, \*\**P* < 0.01). All statistical analyses were conducted using SPSS program (SPSS Inc., Chicago, IL, USA).

## Acknowledgments

This study was supported by the National Research Foundation of Korea (NRF), the Ministry of Education (2017R1A6A1A03015642), Korea.

## Author Contributions

Young-Su Yi, Han Gyung Kim, Ji Hye Kim, and Jae Youl Cho conceived and designed the experiments; Young-Su Yi, Han Gyung Kim, Ji Hye Kim, Woo Seok Yang, Eunji Kim, Deok Jeong, Jae Gwang Park, and Nur Aziz performed the experiments; Young-Su Yi, Han Gyung Kim, Ji Hye Kim, Suk Kim, Narayana Parameswaran, and Jae Youl Cho analyzed the data; Suk Kim and Narayana Parameswaran contributed reagents/materials/analysis tools; Young-Su Yi, Han Gyung Kim, Ji Hye Kim, and Jae Youl Cho wrote the paper.

## Conflicts of Interest

The authors declare no conflicts of interest.

## Supplementary information

### Supplementary Figure Legends

**Figure S1.**
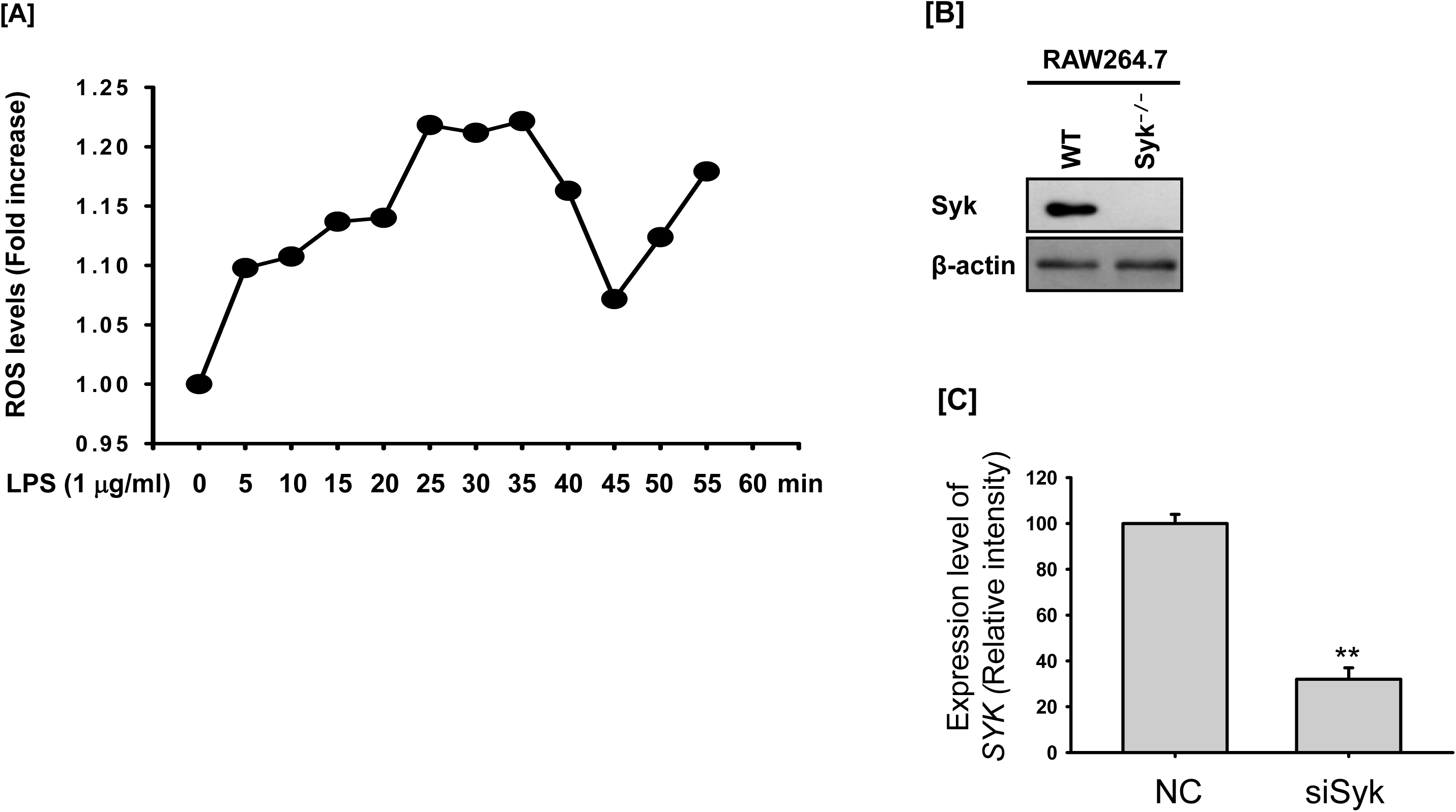
LPS-induced ROS generation in RAW264.7 cells, and the generation of Syk^-/-^ RAW264.7 cells. (A) RAW264.7 cells treated with LPS (1 μg/ml) for the indicated time were incubated with DHR 123 (20 μM) for 20 min, and the ROS level was determined by measuring fluorescence. (B) Syk^-/-^ RAW264.7 cells were generated by the CRISPR/Cas9 system, and the expression of Syk was detected by Western blot analysis. (C) The mRNA level of *SYK* from RAW264.7 cells transfected with siRNAs to NC or Syk was assessed by real-time PCR. N.C.: Negative control.

**Figure S2.**
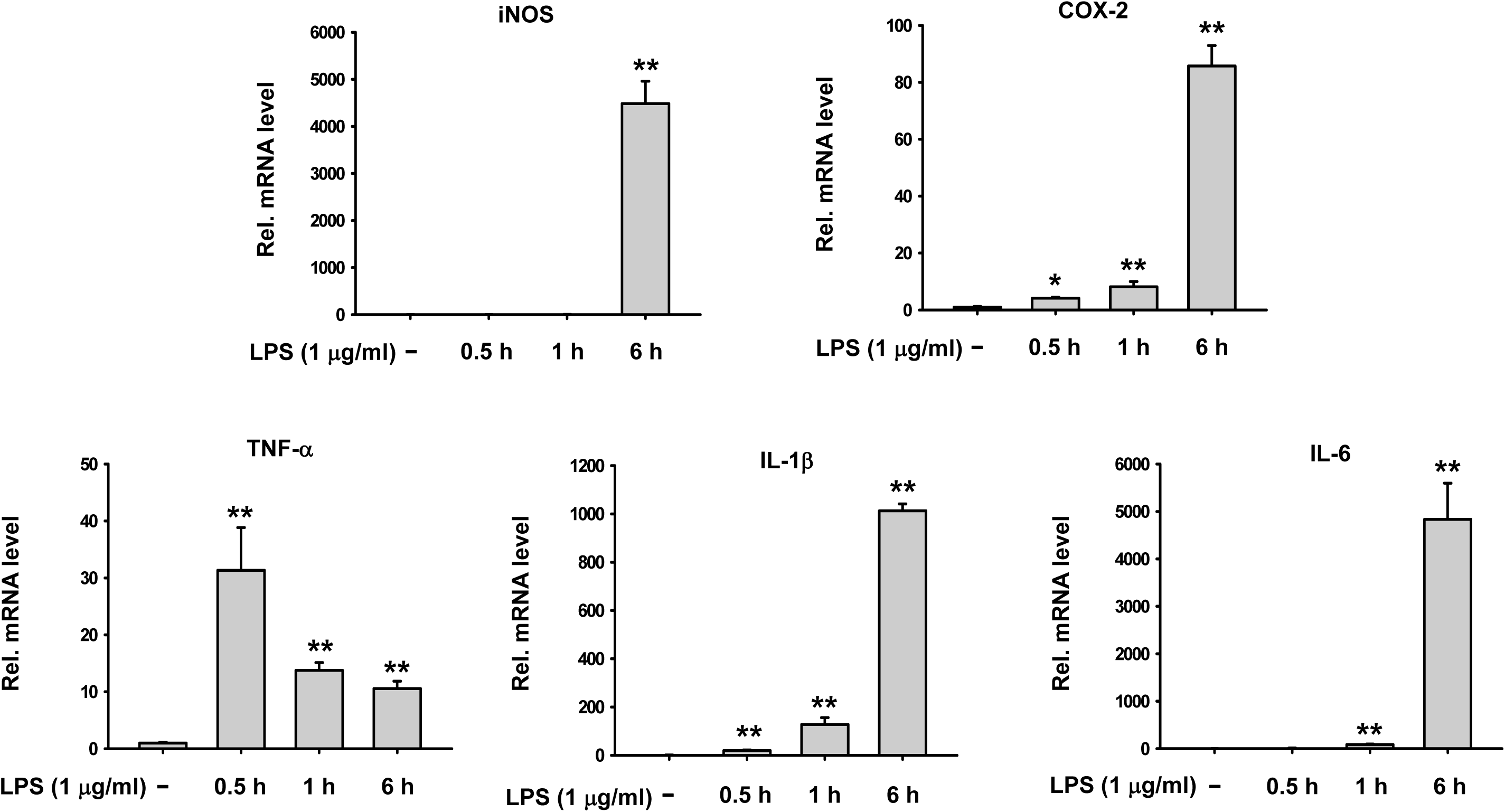
LPS-induced expression of inflammatory genes in RAW264.7 cells. RAW264.7 cells were treated with LPS (1 μg/ml) for the indicated time, and mRNA expression of the target genes was determined by quantitative real time PCR. \**P* < 0.05, \*\**P* < 0.01 compared to controls.

**Figure S3.**
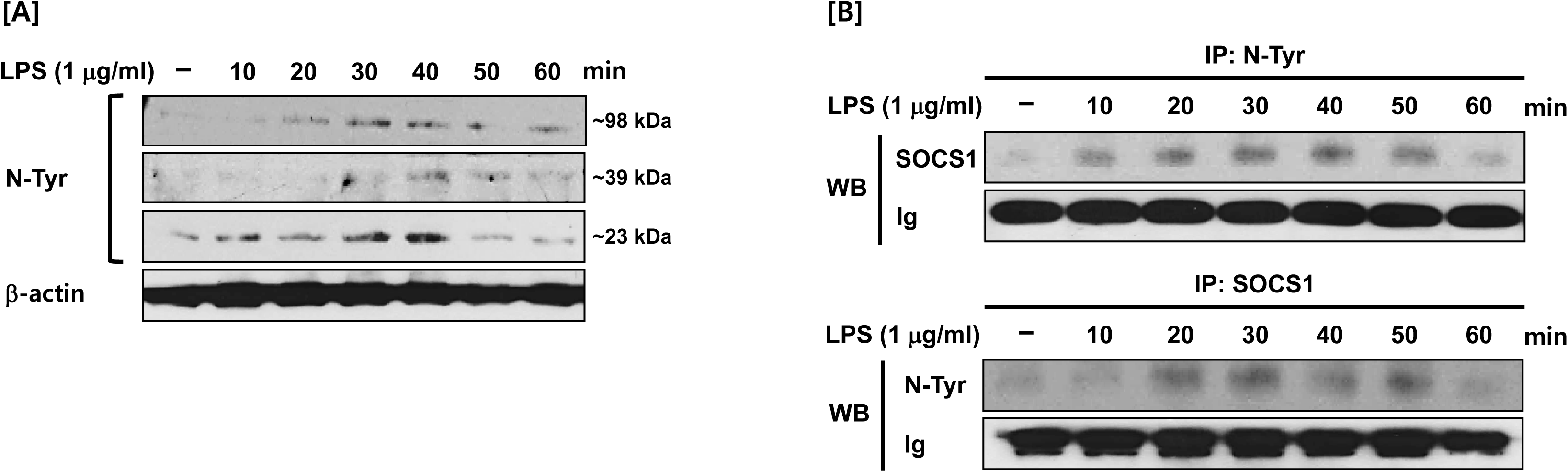
LPS-induced tyrosine nitration on SOCS1 in RAW264.7 cells. (A) N-Tyr proteins in the whole cell lysates of RAW264.7 cells treated with LPS (1 μg/ml) for the indicated time were detected by Western blot analysis. (B) N-Tyr SOCS1 in the whole cell lysates of RAW264.7 cells treated with LPS (1 μg/ml) for the indicated time was detected by immunoprecipitation and Western blot analysis.

**Figure S4.**
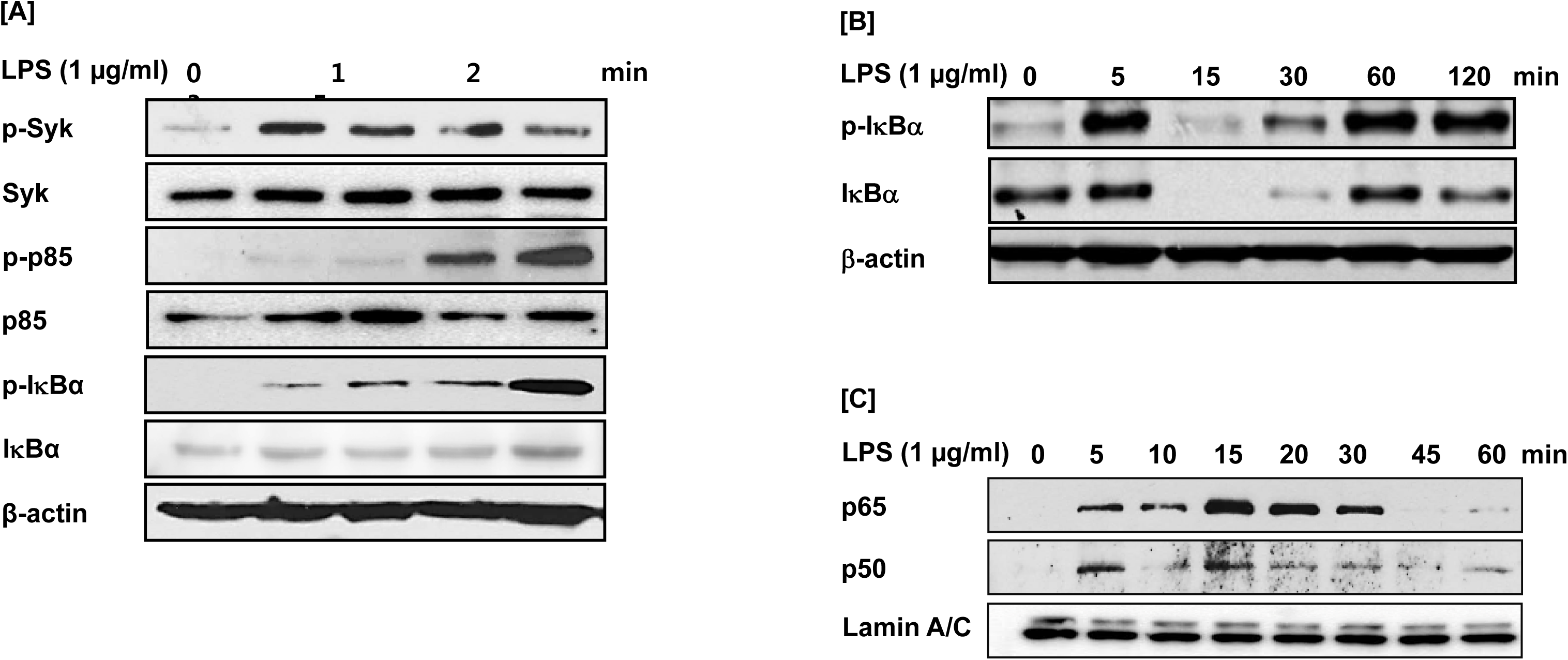
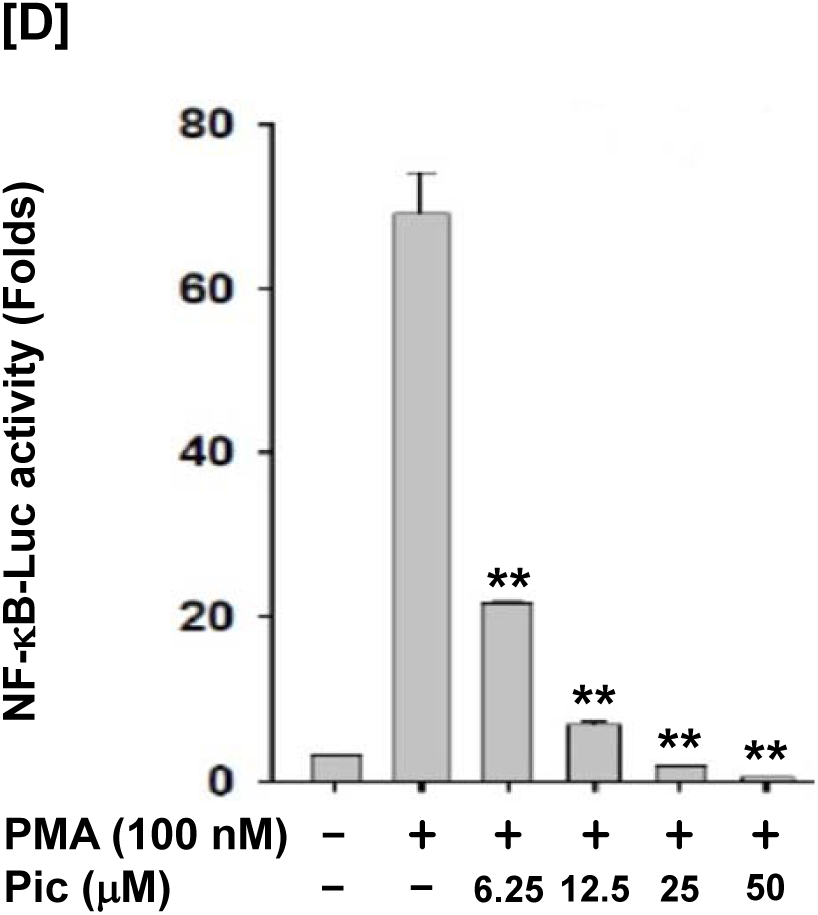
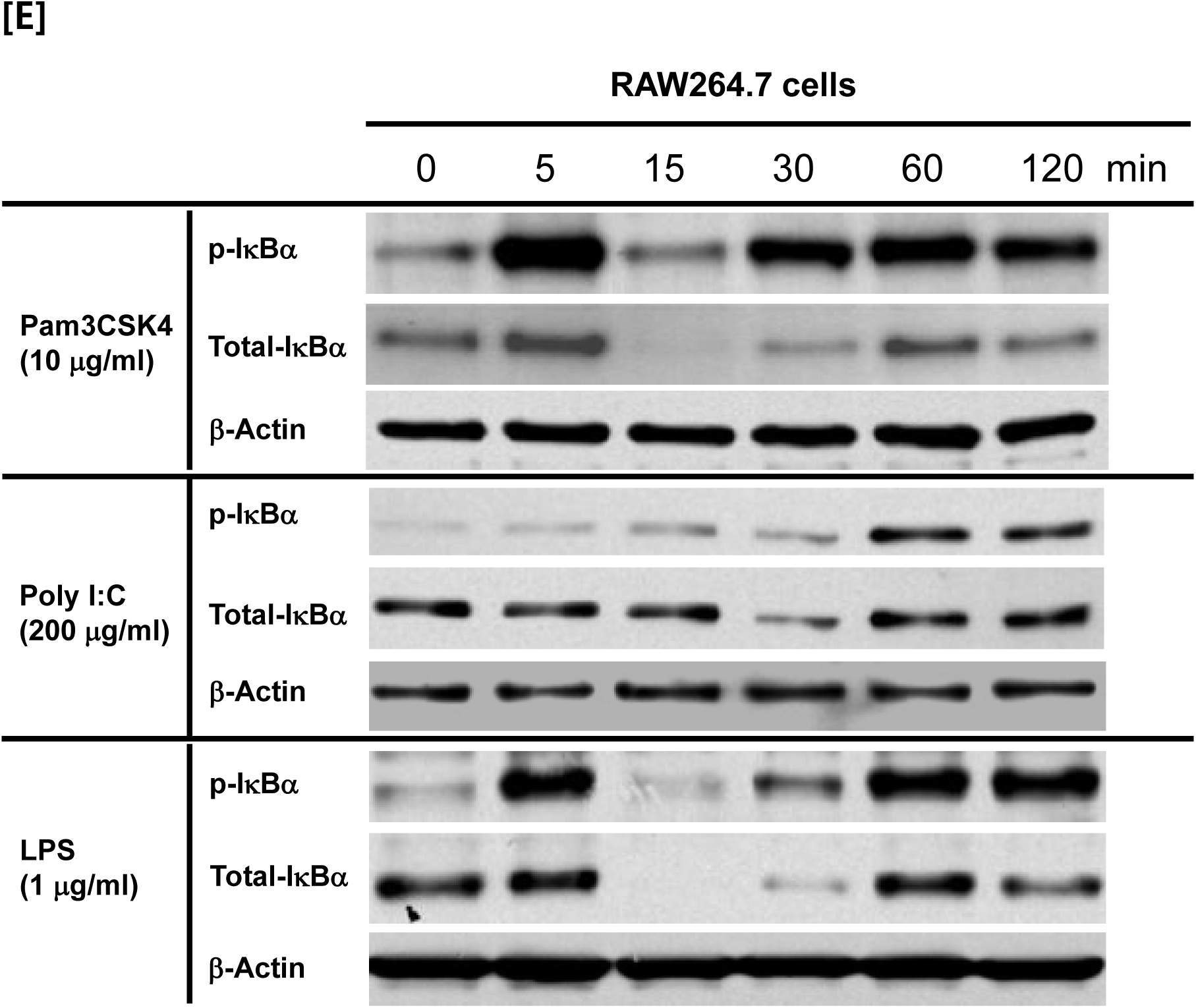
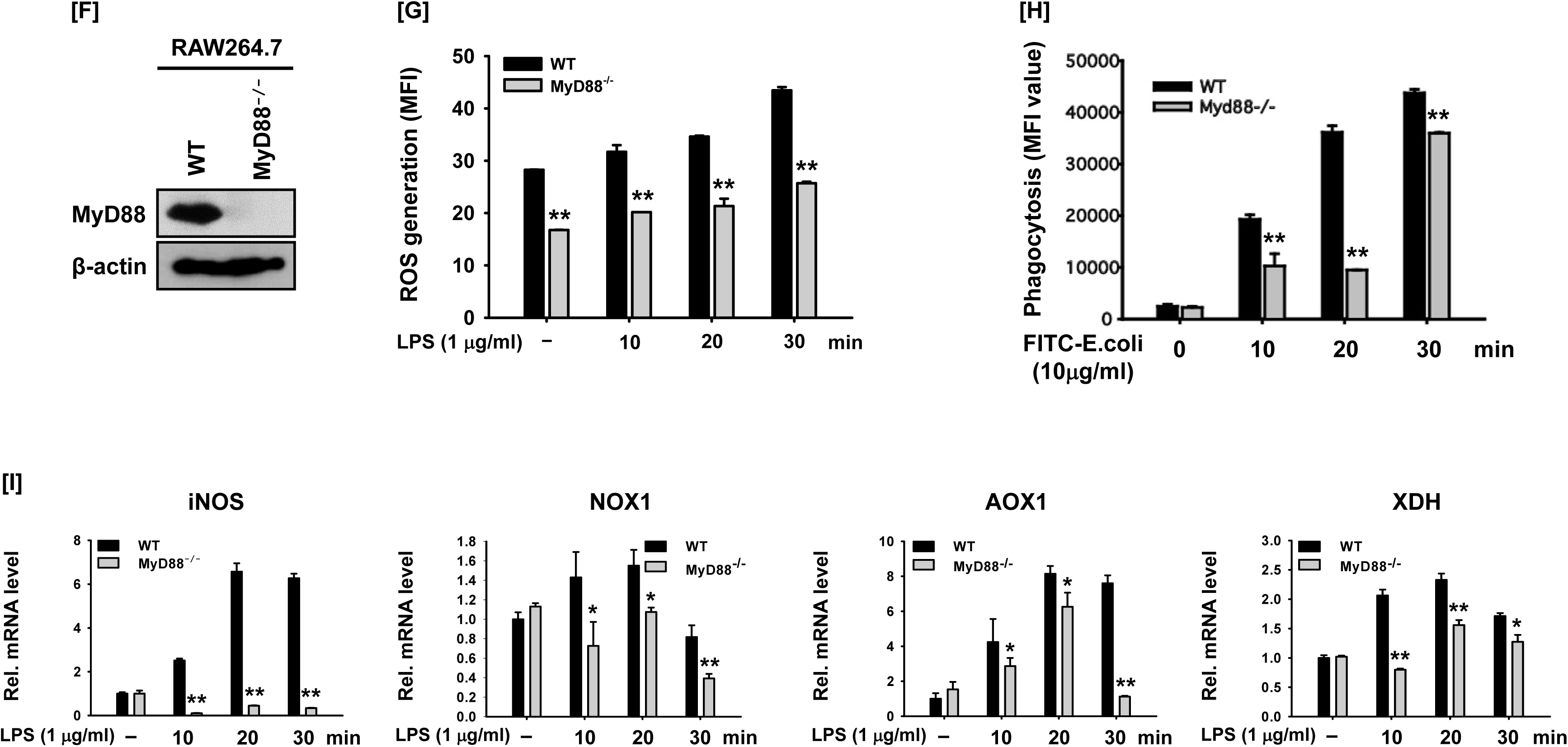
The involvement of Syk on MyD88-dependent NF-kB activation. (A and B) Syk, p-Syk, p85, p-p85, IκBα, and p-IκBα in the whole cell lysates of RAW264.7 cells treated with LPS (1 μg/ml) for the indicated time were detected by Western blot analysis. (C) p65 and p50 in the nuclear lysates of RAW264.7 cells treated with LPS (1 μg/ml) for the indicated time were detected by Western blot analysis. (D) HEK293 cells transfected with the NF-κB-Luciferase reporter gene plasmid were treated with PMA (100 nM) and Pic (0 – 50 μM), and the luciferase activity was measured by a luminometer and normalized to β-galactosidase activity. (E) IκBα and p-IκBα were detected by Western blot analysis in the whole cell lysates of RAW264.7 cells treated with Pam3CSK4 (10 μg/ml), Poly I:C (200 μg/ml), and LPS (1 μg/ml) for the indicated time. (F) MyD88^-/-^ RAW264.7 cells were generated by the CRISPR/Cas9 system, and the expression of MyD88 was detected by Western blot analysis. (G) WT RAW264.7 and MyD88^-/-^ RAW264.7 cells treated with LPS (1 μg/ml) for the indicated time were incubated with DHR 123 (20 μM) for 20 min, and the ROS level was determined by measuring fluorescence. (H) WT RAW264.7 and MyD88^-/-^ RAW264.7 cells were incubated with FITC-E.coli (10 μg/ml) for the indicated time, and the fluorescence was determined by a fluorescence plate reader. (I) WT RAW264.7 and MyD88^-/-^ RAW264.7 cells were treated with LPS (1 μg/ml) for the indicated time, and mRNA expression of the target genes was determined by quantitative real time PCR. \**P* < 0.05, \*\**P* < 0.01 compared to controls.

**Figure S5.**
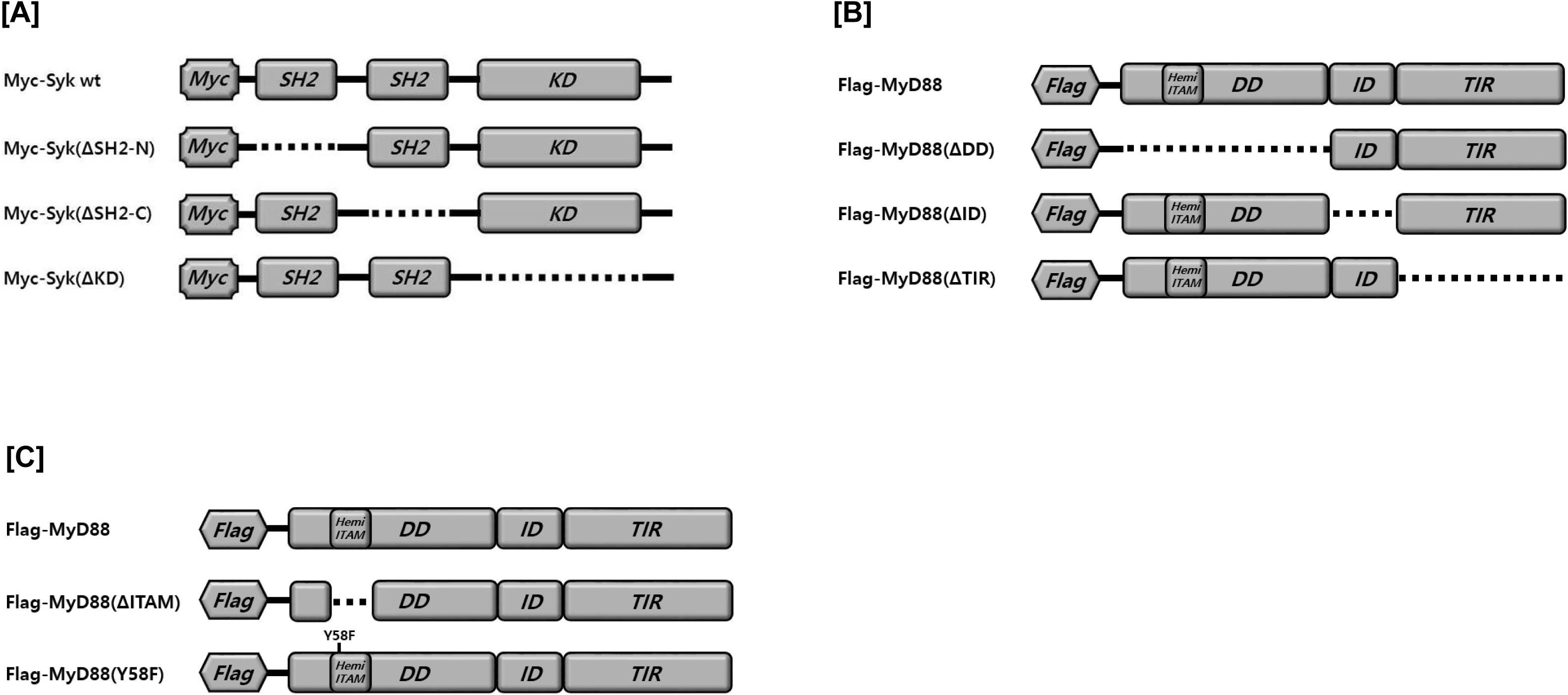
Generation of MyD88 and Syk mutants. Schematic structures of (A) WT and domain deletion mutants of MyD88, (B) WT and domain deletion mutants of Syk, and (C) WT, hemi-ITAM deletion mutant, and Y58F mutant of MyD88.

**Figure S6.**
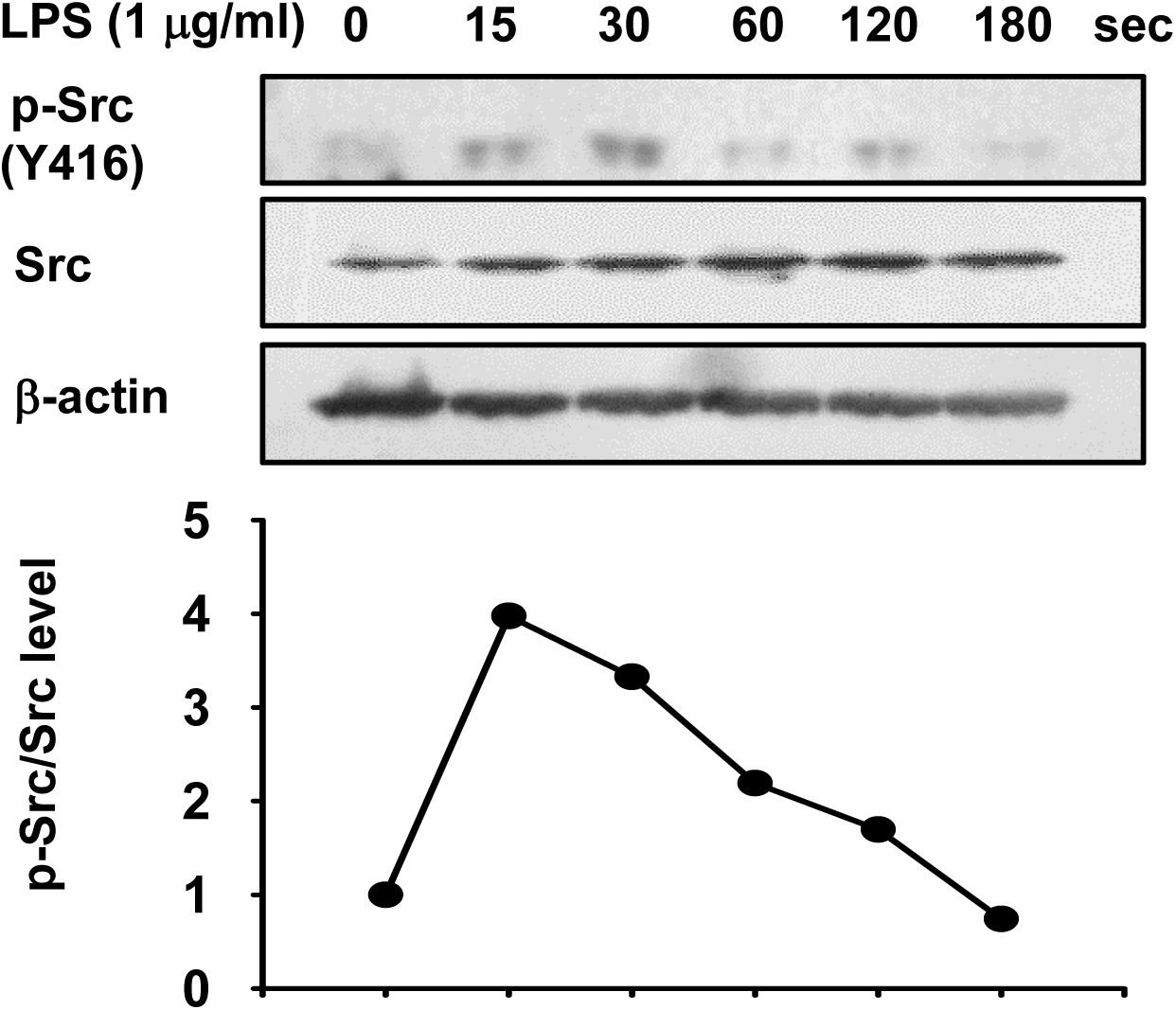
Time-dependent Src phosphorylation profile induced by LPS in RAW264.7 cells. Src and p-Src in the whole cell lysates of RAW264.7 cells treated with LPS (1 μg/ml) for the indicated time were detected by Western blot analysis, and the p-Src/Src level was plotted.

**Figure S7.**
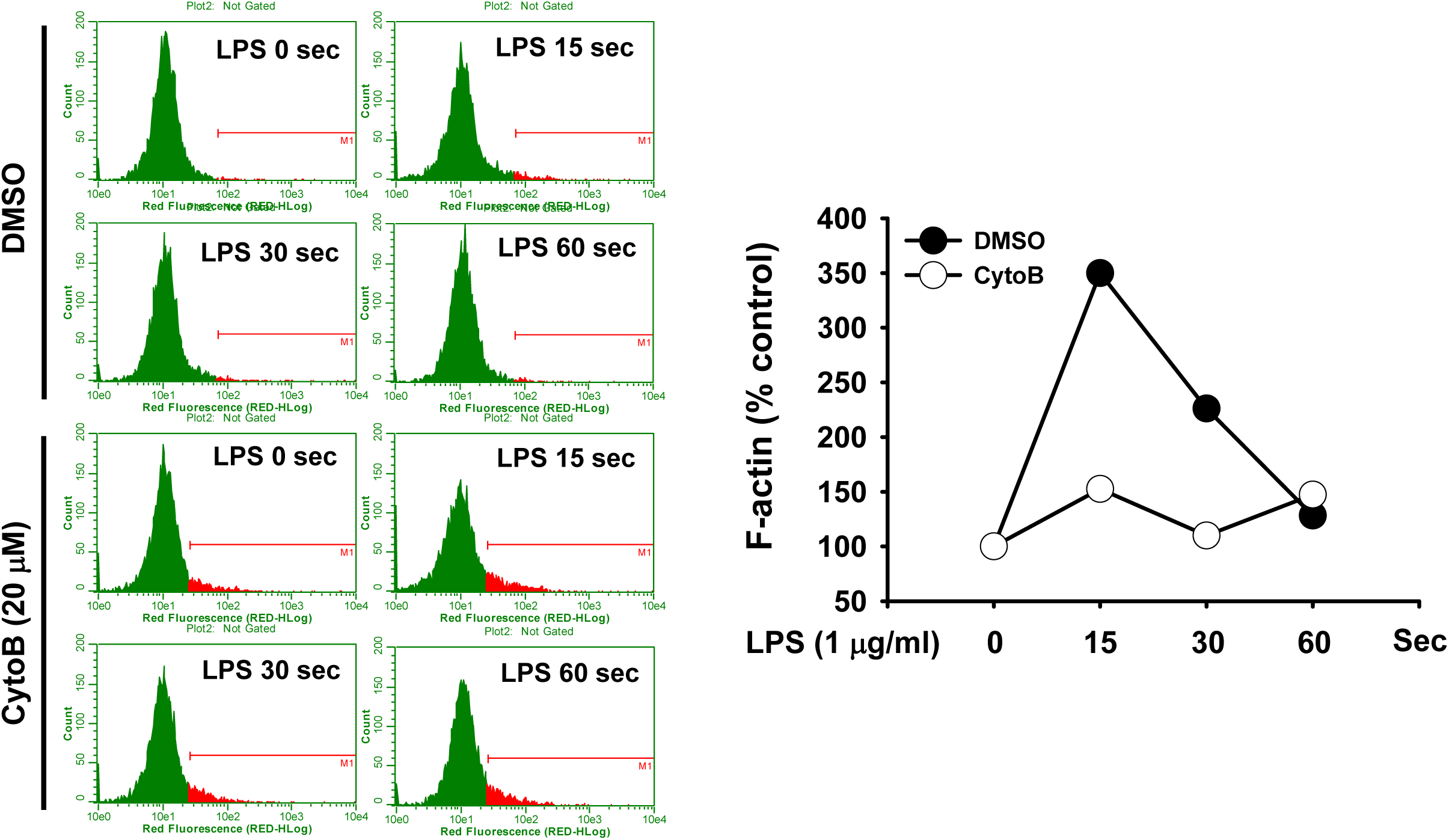
LPS-induced formation of F-actin and the inhibition of LPS-induced F-actin formation by CytoB in RAW264.7 cells. RAW264.7 cells incubated with vehicle (DMSO) or CytoB (20 μM) were treated with LPS (1 μg/ml) for the indicated time in the presence of Texas Red-X phalloidin (200 units/ml), and the fluorescence was determined by a flow cytometer and plotted.

**Supplementary Table 1.**
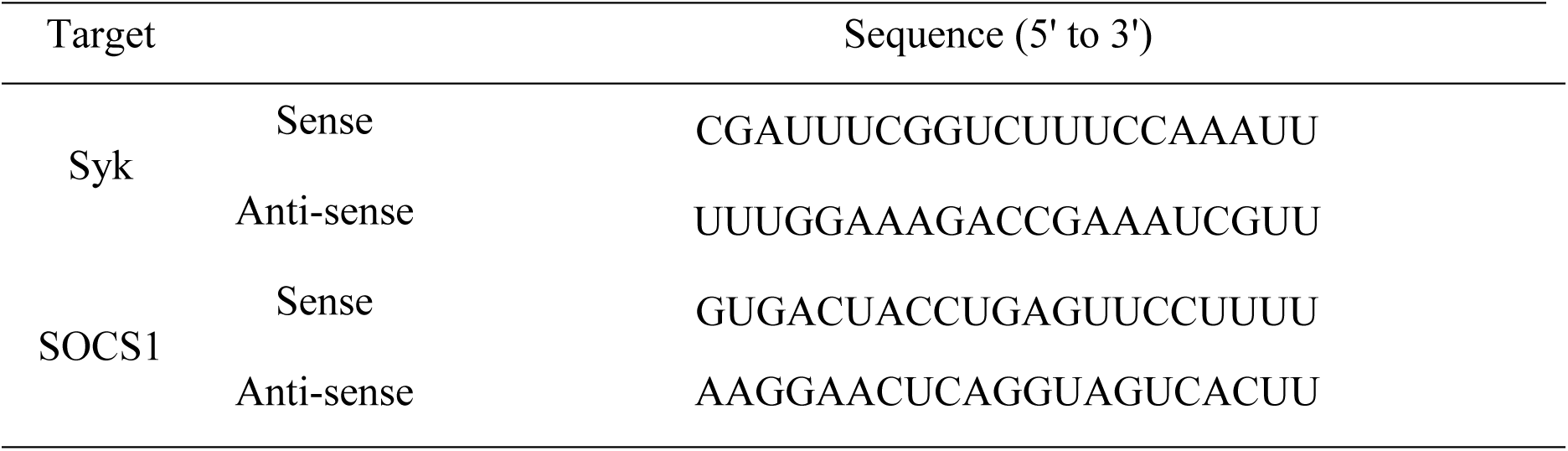
The sequences of siRNAs used in this study

**Supplementary Table 2.**
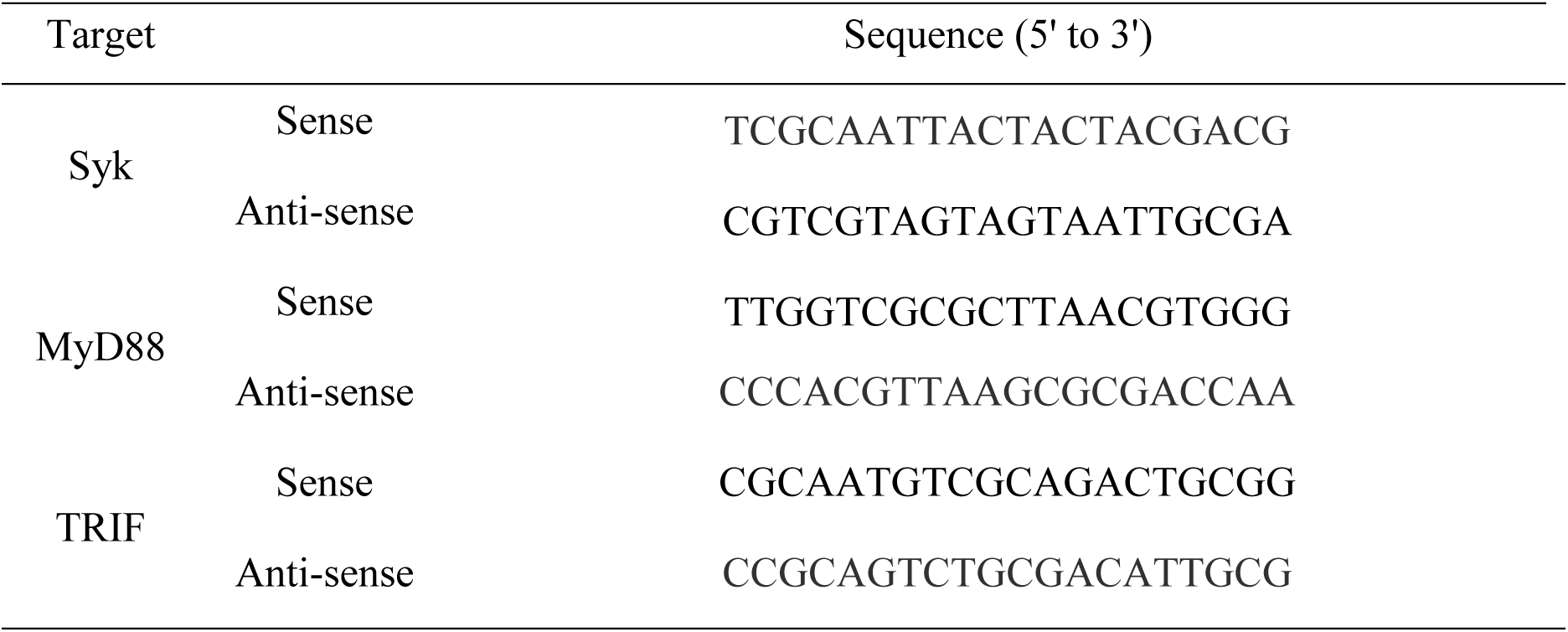
The sequences of sgRNAs used in this study

**Supplementary Table 3.**
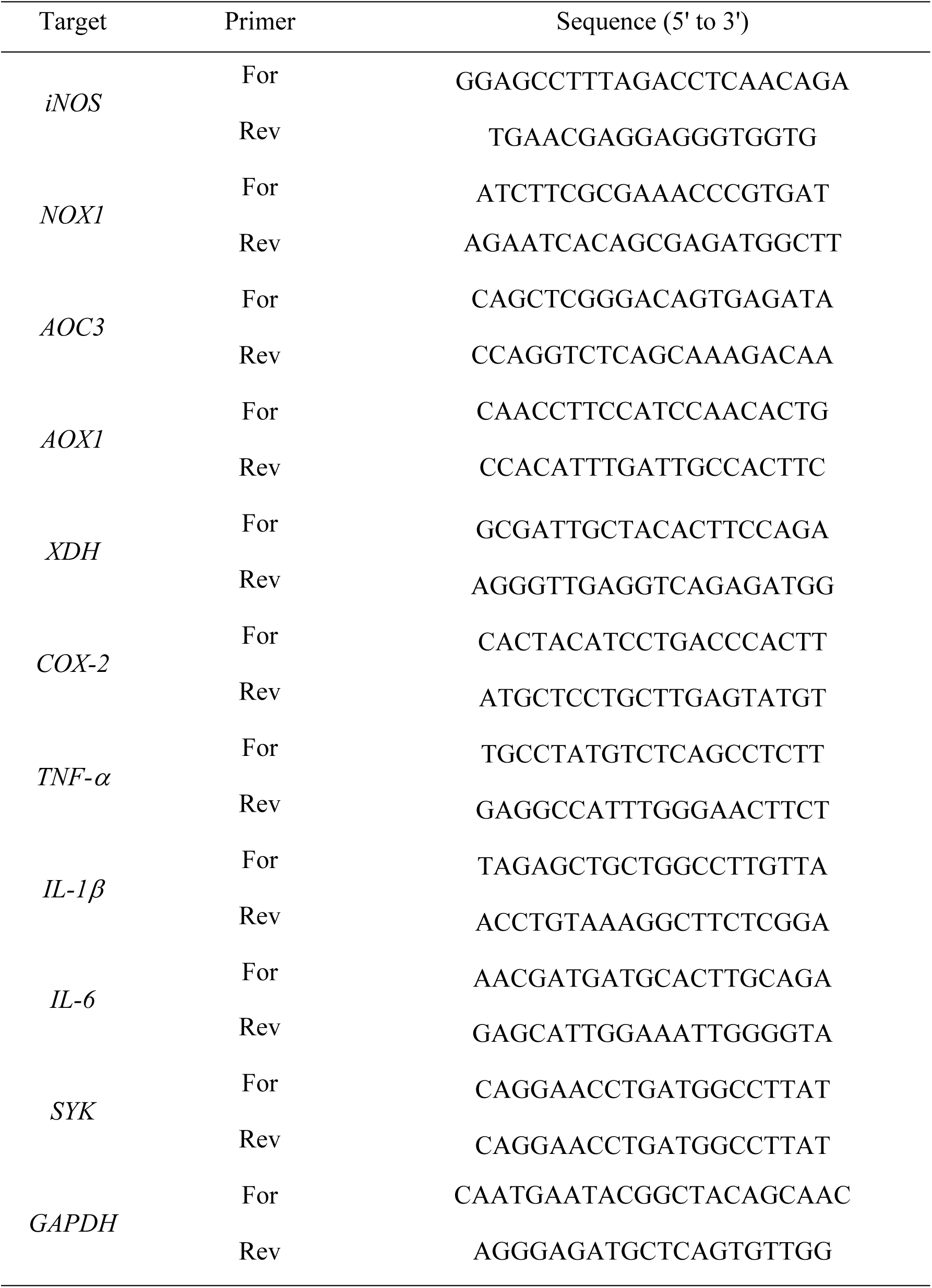
Primer sequences used for quantitative real time PCR in this study

## References

Abram CL, Lowell CA (2007) The expanding role for ITAM-based signaling pathways in immune cells. Sci STKE 2007: re2

Abriata LA, Cassina A, Tortora V, Marin M, Souza JM, Castro L, Vila AJ, Radi R (2009) Nitration of solvent-exposed tyrosine 74 on cytochrome c triggers heme iron-methionine 80 bond disruption. Nuclear magnetic resonance and optical spectroscopy studies. J Biol Chem 284: 17–26

Bezbradica JS, Medzhitov R (2012) Role of ITAM signaling module in signal integration. Curr Opin Immunol 24: 58–66

Bid HK, Roberts RD, Manchanda PK, Houghton PJ (2013) RAC1: an emerging therapeutic option for targeting cancer angiogenesis and metastasis. Mol Cancer Ther 12: 1925–1934

Byeon SE, Yi YS, Oh J, Yoo BC, Hong S, Cho JY (2012) The role of Src kinase in macrophage-mediated inflammatory responses. Mediators of inflammation 2012: 512926

Cao Z, Xiong J, Takeuchi M, Kurama T, Goeddel DV (1996) TRAF6 is a signal transducer for interleukin-1. Nature 383: 443–446

Castro GD, Delgado de Layno AM, Costantini MH, Castro JA (2001) Cytosolic xanthine oxidoreductase mediated bioactivation of ethanol to acetaldehyde and free radicals in rat breast tissue. Its potential role in alcohol-promoted mammary cancer. Toxicology 160: 11–18

Cheng PW, Lee HC, Lu PJ, Chen HH, Lai CC, Sun GC, Yeh TC, Hsiao M, Lin YT, Liu CP, Tseng CJ (2016) Resveratrol Inhibition of Rac1-Derived Reactive Oxygen Species by AMPK Decreases Blood Pressure in a Fructose-Induced Rat Model of Hypertension. Sci Rep 6: 25342

Covarrubias A, Byles V, Horng T (2013) ROS sets the stage for macrophage differentiation. Cell research 23: 984–985

Crusz SM, Balkwill FR (2015) Inflammation and cancer: advances and new agents. Nature reviews Clinical oncology 12: 584–596

Cuadrado A, Martin-Moldes Z, Ye J, Lastres-Becker I (2014) Transcription factors NRF2 and NF-kappaB are coordinated effectors of the Rho family, GTP-binding protein RAC1 during inflammation. J Biol Chem 289: 15244–15258

Cuda CM, Misharin AV, Gierut AK, Saber R, Haines GK, 3rd, Hutcheson J, Hedrick SM, Mohan C, Budinger GS, Stehlik C, Perlman H (2014) Caspase-8 acts as a molecular rheostat to limit RIPK1- and MyD88-mediated dendritic cell activation. J Immunol 192: 5548–5560

de Souza LF,Barreto F, da Silva EG, Andrades ME, Guimaraes EL, Behr GA, Moreira JC, Bernard EA (2007) Regulation of LPS stimulated ROS production in peritoneal macrophages from alloxan-induced diabetic rats: involvement of high glucose and PPARgamma. Life Sci 81: 153–159

Deguine J, Barton GM (2014) MyD88: a central player in innate immune signaling. F1000prime reports 6: 97

Du L, Zhou J, Zhang J, Yan M, Gong L, Liu X, Chen M, Tao K, Luo N, Liu J (2012) Actin filament reorganization is a key step in lung inflammation induced by systemic inflammatory response syndrome. Am J Respir Cell Mol Biol 47: 597–603

Hall A (1998) Rho GTPases and the actin cytoskeleton. Science 279: 509–514

Holtorf A, Conrad A, Holzmann B, Janssen KP (2018) Cell-type specific MyD88 signaling is required for intestinal tumor initiation and progression to malignancy. Oncoimmunology 7: e1466770

Janeway CA, Jr., Medzhitov R (2002) Innate immune recognition. Annual review of immunology 20: 197–216

Janssens S, Burns K, Tschopp J, Beyaert R (2002) Regulation of interleukin-1- and lipopolysaccharide-induced NF-kappaB activation by alternative splicing of MyD88. Current biology : CB 12: 467–471

Janssens S, Burns K, Vercammen E, Tschopp J, Beyaert R (2003) MyD88S, a splice variant of MyD88, differentially modulates NF-kappaB- and AP-1-dependent gene expression. FEBS letters 548: 103–107

Jerkic M, Sotov V, Letarte M (2012) Oxidative stress contributes to endothelial dysfunction in mouse models of hereditary hemorrhagic telangiectasia. Oxid Med Cell Longev 2012: 686972

Kim HG, Choi S, Lee J, Hong YH, Jeong D, Yoon K, Yoon DH, Sung GH, Lee S, Hong S, Yi YS, Kim JH, Cho JY (2018) Src Is a Prime Target Inhibited by Celtis choseniana Methanol Extract in Its Anti-Inflammatory Action. Evidence-based complementary and alternative medicine : eCAM 2018: 3909038

Kim HG, Yang WS, Hong YH, Kweon DH, Lee J, Kim S, Cho JY (2019) Anti-inflammatory functions of the CDC25 phosphatase inhibitor BN82002 via targeting AKT2. Biochemical pharmacology 164: 216–227

Kim JH, Kim SC, Yi YS, Yang WS, Yang Y, Kim HG, Lee JY, Kim KH, Yoo BC, Hong S, Cho JY (2013) Adenosine dialdehyde suppresses MMP-9-mediated invasion of cancer cells by blocking the Ras/Raf-1/ERK/AP-1 signaling pathway. Biochemical pharmacology 86: 1285–1300

Kundu TK, Hille R, Velayutham M, Zweier JL (2007) Characterization of superoxide production from aldehyde oxidase: an important source of oxidants in biological tissues. Archives of biochemistry and biophysics 460: 113–121

Lee YG, Chain BM, Cho JY (2009) Distinct role of spleen tyrosine kinase in the early phosphorylation of inhibitor of kappaB alpha via activation of the phosphoinositide-3-kinase and Akt pathways. The international journal of biochemistry & cell biology 41: 811–821

Lim JE, Song M, Jin J, Kou J, Pattanayak A, Lalonde R, Fukuchi K (2012) The effects of MyD88 deficiency on exploratory activity, anxiety, motor coordination, and spatial learning in C57BL/6 and APPswe/PS1dE9 mice. Behavioural brain research 227: 36–42

Lin YT, Wu YC, Sun GC, Ho CY, Wong TY, Lin CH, Chen HH, Yeh TC, Li CJ, Tseng CJ, Cheng PW (2018) Effect of Resveratrol on Reactive Oxygen Species-Induced Cognitive Impairment in Rats with Angiotensin II-Induced Early Alzheimer’s Disease (dagger). J Clin Med 7

Liu HT, Du YG, He JL, Chen WJ, Li WM, Yang Z, Wang YX, Yu C (2010) Tetramethylpyrazine inhibits production of nitric oxide and inducible nitric oxide synthase in lipopolysaccharide-induced N9 microglial cells through blockade of MAPK and PI3K/Akt signaling pathways, and suppression of intracellular reactive oxygen species. Journal of ethnopharmacology 129: 335–343

Lucas K, Maes M (2013) Role of the Toll Like receptor (TLR) radical cycle in chronic inflammation: possible treatments targeting the TLR4 pathway. Molecular neurobiology 48: 190–204

Lyles GA (1996) Mammalian plasma and tissue-bound semicarbazide-sensitive amine oxidases: biochemical, pharmacological and toxicological aspects. The international journal of biochemistry & cell biology 28: 259–274

MacLean-Fletcher S, Pollard TD (1980) Mechanism of action of cytochalasin B on actin. Cell 20: 329–341

Makhezer N, Ben Khemis M, Liu D, Khichane Y, Marzaioli V, Tlili A, Mojallali M, Pintard C, Letteron P, Hurtado-Nedelec M, El-Benna J, Marie JC, Sannier A, Pelletier AL, Dang PM (2019) NOX1-derived ROS drive the expression of Lipocalin-2 in colonic epithelial cells in inflammatory conditions. Mucosal immunology 12: 117–131

Mocsai A, Ruland J, Tybulewicz VL (2010) The SYK tyrosine kinase: a crucial player in diverse biological functions. Nature reviews Immunology 10: 387–402

Mohs A, Kuttkat N, Otto T, Youssef SA, De Bruin A, Trautwein C (2019) MyD88-dependent signaling in non-parenchymal cells promotes liver carcinogenesis. Carcinogenesis

Nagase M, Fujita T (2013) Role of Rac1-mineralocorticoid-receptor signalling in renal and cardiac disease. Nat Rev Nephrol 9: 86–98

Oda K, Kitano H (2006) A comprehensive map of the toll-like receptor signaling network. Molecular systems biology 2: 2006 0015

Park JG, Son YJ, Yoo BC, Yang WS, Kim JH, Cho JY (2017) Syk Plays a Critical Role in the Expression and Activation of IRAK1 in LPS-Treated Macrophages. Mediators of inflammation 2017: 1506248

Rowley RB, Bolen JB, Fargnoli J (1995) Molecular cloning of rodent p72Syk. Evidence of alternative mRNA splicing. The Journal of biological chemistry 270: 12659–12664

Seow CJ, Chue SC, Wong WS (2002) Piceatannol, a Syk-selective tyrosine kinase inhibitor, attenuated antigen challenge of guinea pig airways in vitro. Eur J Pharmacol 443: 189–196

Shalem O, Sanjana NE, Hartenian E, Shi X, Scott DA, Mikkelson T, Heckl D, Ebert BL, Root DE, Doench JG, Zhang F (2014) Genome-scale CRISPR-Cas9 knockout screening in human cells. Science 343: 84–87

Siracusano G, Venuti A, Lombardo D, Mastino A, Esclatine A, Sciortino MT (2016) Early activation of MyD88-mediated autophagy sustains HSV-1 replication in human monocytic THP-1 cells. Scientific reports 6: 31302

Steele RE, Stover NA, Sakaguchi M (1999) Appearance and disappearance of Syk family protein-tyrosine kinase genes during metazoan evolution. Gene 239: 91–97

Suschek CV, Schnorr O, Kolb-Bachofen V (2004) The role of iNOS in chronic inflammatory processes in vivo: is it damage-promoting, protective, or active at all? Current molecular medicine 4: 763–775

Takeda K, Akira S (2005) Toll-like receptors in innate immunity. Int Immunol 17: 1–14

Takeuchi O, Akira S (2010) Pattern recognition receptors and inflammation. Cell 140: 805–820

Taniguchi T, Kobayashi T, Kondo J, Takahashi K, Nakamura H, Suzuki J, Nagai K, Yamada T, Nakamura S, Yamamura H (1991) Molecular cloning of a porcine gene syk that encodes a 72-kDa protein-tyrosine kinase showing high susceptibility to proteolysis. The Journal of biological chemistry 266: 15790–15796

Tata A, Stoppel DC, Hong S, Ben-Zvi A, Xie T, Gu C (2014) An image-based RNAi screen identifies SH3BP1 as a key effector of Semaphorin 3E-PlexinD1 signaling. J Cell Biol 205: 573–590

Thews O, Lambert C, Kelleher DK, Biesalski HK, Vaupel P, Frank J (2005) Possible protective effects of alpha-tocopherol on enhanced induction of reactive oxygen species by 2-methoxyestradiol in tumors. Adv Exp Med Biol 566: 349–355

Todoric J, Antonucci L, Karin M (2016) Targeting Inflammation in Cancer Prevention and Therapy. Cancer Prev Res (Phila) 9: 895–905

van der Vaart M, Korbee CJ, Lamers GE, Tengeler AC, Hosseini R, Haks MC, Ottenhoff TH, Spaink HP, Meijer AH (2014) The DNA damage-regulated autophagy modulator DRAM1 links mycobacterial recognition via TLR-MYD88 to autophagic defense [corrected]. Cell host & microbe 15: 753–767

Wu CM, Cheng YL, Dai YH, Chen MF, Wang CC (2014) alpha-Tocopherol protects keratinocytes against ultraviolet A irradiation by suppressing glutathione depletion, lipid peroxidation and reactive oxygen species generation. Biomed Rep 2: 419–423

Xue L, Geahlen RL, Tao WA (2013) Identification of direct tyrosine kinase substrates based on protein kinase assay-linked phosphoproteomics. Molecular & cellular proteomics : MCP 12: 2969–2980

Yamakura F, Taka H, Fujimura T, Murayama K (1998) Inactivation of human manganese-superoxide dismutase by peroxynitrite is caused by exclusive nitration of tyrosine 34 to 3-nitrotyrosine. J Biol Chem 273: 14085–14089

Yang Y, Kim SC, Yu T, Yi YS, Rhee MH, Sung GH, Yoo BC, Cho JY (2014) Functional roles of p38 mitogen-activated protein kinase in macrophage-mediated inflammatory responses. Mediators of inflammation 2014: 352371

Yi YS (2016a) Folate Receptor-Targeted Diagnostics and Therapeutics for Inflammatory Diseases. Immune network 16: 337–343

Yi YS (2016b) Functional Role of Milk Fat Globule-Epidermal Growth Factor VIII in Macrophage-Mediated Inflammatory Responses and Inflammatory/Autoimmune Diseases. Mediators of inflammation 2016: 5628486

Yi YS (2017) Caspase-11 non-canonical inflammasome: a critical sensor of intracellular lipopolysaccharide in macrophage-mediated inflammatory responses. Immunology 152: 207–217

Yi YS (2018a) Regulatory Roles of the Caspase-11 Non-Canonical Inflammasome in Inflammatory Diseases. Immune network 18: e41

Yi YS (2018b) Role of inflammasomes in inflammatory autoimmune rheumatic diseases. The Korean journal of physiology & pharmacology : official journal of the Korean Physiological Society and the Korean Society of Pharmacology 22: 1–15

Yi YS, Jian J, Gonzalez-Gugel E, Shi YX, Tian Q, Fu W, Hettinghouse A, Song W, Liu R, He M, Qi H, Yang J, Du X, Xiao G, Chen L, Liu CJ (2018) p204 Is Required for Canonical Lipopolysaccharide-induced TLR4 Signaling in Mice. EBioMedicine 29: 78–91

Yi YS, Son YJ, Ryou C, Sung GH, Kim JH, Cho JY (2014) Functional roles of Syk in macrophage-mediated inflammatory responses. Mediators of inflammation 2014: 270302

Yu T, Yi YS, Yang Y, Oh J, Jeong D, Cho JY (2012) The pivotal role of TBK1 in inflammatory responses mediated by macrophages. Mediators of inflammation 2012: 979105

Ziegenfuss JS, Biswas R, Avery MA, Hong K, Sheehan AE, Yeung YG, Stanley ER, Freeman MR (2008) Draper-dependent glial phagocytic activity is mediated by Src and Syk family kinase signalling. Nature 453: 935–939

